# Optical Metabolic Imaging Uncovers Sex- and Diet-dependent Lipid Changes in Aging Drosophila Brain

**DOI:** 10.1101/2022.10.01.510416

**Authors:** Yajuan Li, Phyllis Chang, Shiriya Sankaran, Hongje Jang, Yuhang Nie, Audrey Zeng, Sahran Hussain, Jane Y. Wu, Xu Chen, Lingyan Shi

**Affiliations:** Department of Bioengineering, University of California, San Diego; Department of Neurosciences, School of Medicine, University of California, San Diego; Dept. of Neurology, Feinberg School of Medicine, Northwestern University, Chicago, IL, USA

## Abstract

Aging is associated with progressive declines in physiological integrity and functions alongside increases in vulnerability to develop a number of diseases. The brain regulates sensory and motor functions as well as endocrine functions, and age-associated changes in brain are likely prerequisite for the organismal aging. Lipid metabolism has been associated with brain aging, which could be easily intervened by diets and lifestyles. However, the underlying mechanism through which brain lipid metabolism is regulated by diet during aging is elusive. Using stimulated Raman scattering (SRS) imaging combined with deuterium water (D2O) labeling, we visualized that lipid metabolic activities were changed by diet manipulation in aging *Drosophila* brain. Furthermore, we illuminated that insulin/IGF-1 signaling (IIS) pathway mediates the transformation of brain lipid metabolic changes in both an aging- and a diet-dependent manner. The lipid droplets (LDs) in the brain gradually became inert in both activities of lipid synthesis and mobilization with aging. High sugar diets enhanced the metabolic activity through promoting lipogenesis while dietary restriction increased the metabolic activity in both lipogenesis and lipolysis in brain LDs. However, these effects were impaired in both *chico*^1/+^ and *dfoxo Drosophila* mutants. We also observed that old *chico*^1/+^ brains maintained high metabolic activities, whilst the aged *dfoxo* brains acted exactly the opposite. More interestingly, the sexual dimorphism in brain lipid metabolism was impaired under diet regulation in both *chico*^1/+^ and *dfoxo* mutants. Locally reduced IIS activity in glial cells can mimic the systemic changes in systematic IIS mutants to maintain lipogenesis and lipolysis in aged brains, providing mechanistic insight into the anti-aging effects of IIS pathway. Our results highlight the manipulation of glia-specific IIS activity as a promising strategy in anti-aging treatments.

## Introduction

Aging is an almost universal phenomenon for living organisms and is broadly defined as any change in an organism over time following development. Aging is associated with the progressive deterioration of physiological function and fertility alongside an increased susceptibility to develop a number of diseases and death^1^. Lipid metabolism has been associated with aging and age-related diseases^2^. The brain is a lipid-enriched organ, second only to adipose tissue^3^. Genome-wide association study (GWAS) suggests that age-related changes of lipid metabolism in the brain may be essential for healthy aging and longevity^4^. The brain membrane lipids can be grouped into sphingolipids, glycerophospholipids, and cholesterol^5^. These lipids are used as the building-up materials for cell and organelle membranes, and second messengers for the signaling transductions as well as the energy storage for maintaining cellular functions.

Previous studies observed alterations of lipid metabolism during aging, with a decline of omega-3 fatty acids and an increase in lipid peroxidation^6,7^. A diet deficiency of omega-3 may accelerate neuronal degeneration^8,9^, which indicates brain lipid profile is closely related to dietary nutritional profiles. Another group of lipids that play non-structural roles in the brain are triacylglycerol (TAGs) and steryl esters, which are wrapped in lipid droplets (LDs) and used for energy storage. Study using electron microscopy sections demonstrated that the LDs in nervous system are mainly localized in glial cells instead of nerve cells^10^. Recent research reveals new insights related to the role of LDs in neurodegeneration and normal aging processes. For example, reactive oxygen species (ROS) induced by mitochondrial dysfunction in nerve cells can increase the activities of c-Jun-N-terminal Kinase (JNK) and sterol regulatory element binding protein (SREBP), leading to the accumulation of LDs in glial cells, which occurs when nerve cell apoptosis begins. This demonstrates that the increase of LDs in glial cells is an early transient indicator of nerve cell apoptosis, or it promotes nerve cell death^11^. The inability to transport lipids to glia for LD formation leads to accelerated neurodegeneration under stress^12^. The LDs in glial cells can protect polyunsaturated fatty acids in brain membranes from oxidation under hypoxia and oxidative stress to maintain the proliferation of precursor nerve cells under oxidative stress^13^. The accumulation of LDs in microglia were also described in mouse and human aging brains. The pathological glial cells, named as lipid-droplet-accumulating microglia (LDAM), are defective in phagocytosis, produce high levels of ROS and secrete proinflammatory cytokines. The LDAMs are probably involved in age-related and genetic forms of neurodegenerations^14^.

Despite these efforts, the dynamics of lipid metabolism during brain aging and in different sexes are still unclear. Alterations in nutrition balance is a notable cause of aging^15,16^, but how brain lipid metabolism is modulated by different diets, and the role of Insulin/IGF-1 signaling (IIS) pathway in modulating brain lipid metabolism are still unknown. Addressing these questions will provide insights in studying age-related neurodegenerative diseases.

In this study, we used heavy water (D_2_O)-probed stimulated Raman scattering (DO-SRS) imaging^17,18^ to investigate brain lipid metabolism in female and male *Drosophila* aging brain. We found heterogeneous volumetric distribution of newly synthesized lipids in brain. Sex dimorphism was also displayed in LD metabolism in aged flies. We found lipogenic diets improved brain functions by promoting lipid biosynthesis via the IIS pathway.

## Results

### SRS imaging visualizes lipid metabolic dynamics in *Drosophila* aging brain

We first visualized and analyzed contents of lipids and proteins in young (5-day) and old (35-day) *Drosophila* using Raman spectroscopy. We observed dramatic reduction of total lipids level in aged brains, as indicated by the declined intensity of Raman peak at 2850 cm^-1^ (CH2 from lipids) (Figures 1A and S1A). The peaks at 2930 cm^−1^ (CH3 stretching modes from proteins) remained unchanged during aging. We then quantified changes of lipid relatively to protein by taking the ratios of intensities at 2850 cm^-1^ to those at 2930 cm^-1^, and found significant reduction of lipids in aged flies (Figure 1B)^19^. Unsaturated lipids (peak at 1656 cm^-1^), relatively to protein also showed a significant reduction with aging (Figures S1A-C).

**Figure 1.**
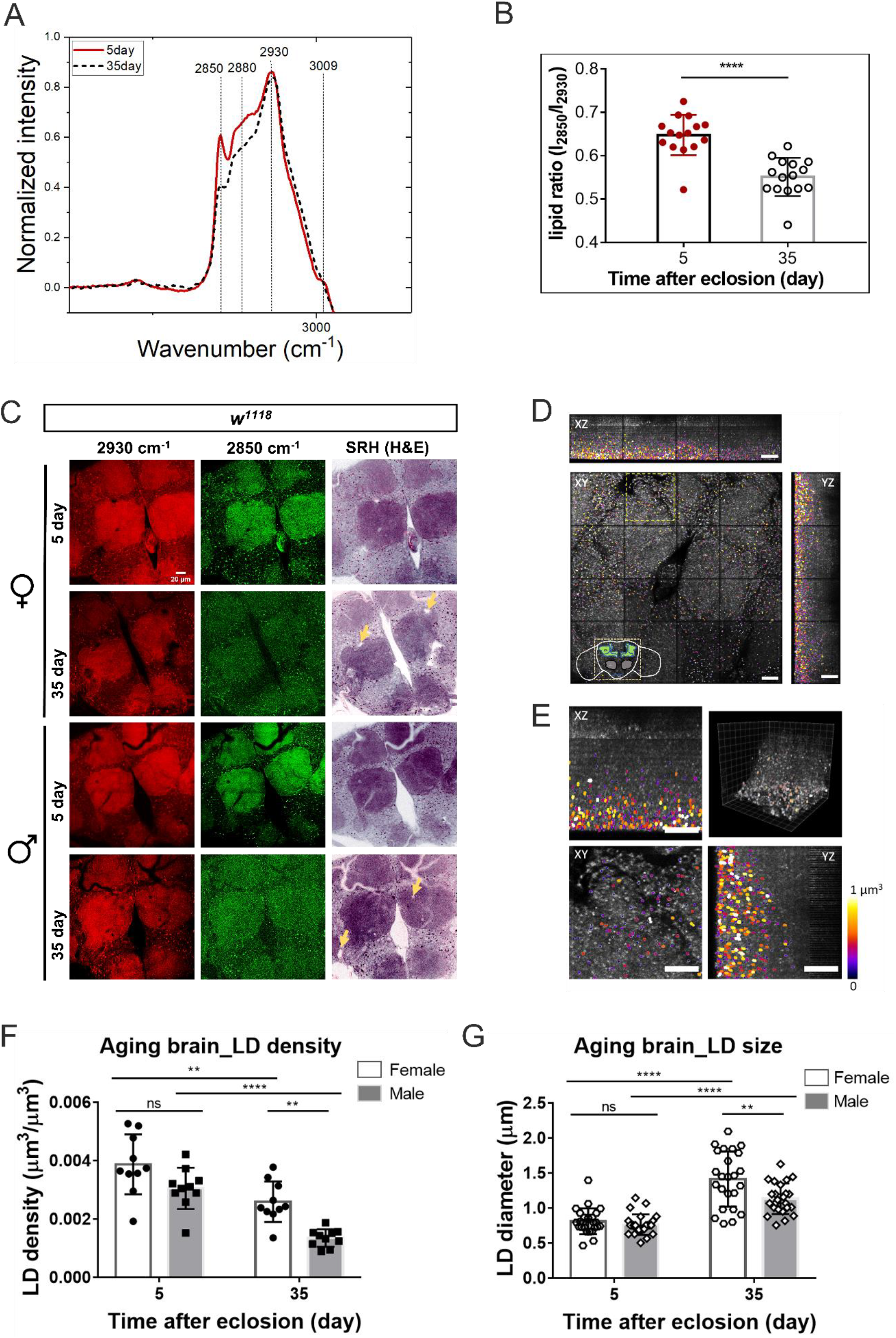
Lipid metabolic changes in *Drosophila* brain are visualized by SRS microscopy. (A) Raman spectra of brains from 5-day and 35-day old flies. The peaks at 2850 cm^-1^, 2880 cm^-1^, 2930 cm^-1^, and 3009 cm^-1^ correspond to lipids, asymmetric CH2 stretching bond, proteins, and unsaturated lipids, respectively. (B) Quantification of the ratios of lipids (2850 cm^-1^) to proteins (2930 cm^-1^). The results were plotted as Mean ± SD. n =15 for each age group. (C) SRS imaging of the central brains from 5-day and 35-day flies. The distributions of proteins and lipids were collected at 2930 cm^-1^ and 2850 cm^-1^ respectively. Stimulated Raman histology (SRH) images were generated from proteins and lipids channels. The orange arrows show the vacuoles in old brains. (D) Volumetric images of lipid metabolism in the entire central brain. Z-stack images were taken at 2850 cm^-1^. (E) Magnified images of boxed regions in (D). (F) and (G) Quantitative comparison of LD density (F) and diameter (G) in both female and male 5-day and 35-day fly brains. Results were plotted as Mean ± SD and compared. N =8∼10 brains for each age group. Statistical significance was determined by student’s *t*-test (B) or 2-way ANOVA (F and G). *, p < 0.05; **, p < 0.01; ***, p < 0.001; ****, p < 0.0001; ns, non-significant difference. Scale bar: 20μm.

We next imaged cellular distributions of proteins and lipids, respectively, in Drosophila brain with SRS microscopy imaging. Raman imaging showed the abundance of total lipids was largely reduced in multiple regions of aged brains, including the central brains (CBs) and optic lobes (OLs), in both sexes (Figures 1C and S1E). The lipid reduction phenotype in aging brains was reported in a number of animal models^20,21^, indicating the same underlying mechanisms in different model systems. In addition, stimulated Raman histology (SRH)^22^, generated based on the proteins (2930 cm^-1^) and lipids (2850 cm^-1^) channels, clearly displayed structural changes during *Drosophila* aging (Figures 1C and S1E). Compared with essentially normal morphology shown in young adults (5-day), gross distortions together with vacuoles and vague outlines of the neuropil (the highly densified compartments of central brains: MBs and ALs) were developed in aged (35-day) flies.

We visualized single LDs using SRS imaging at 2850 cm^-1^ in adult brains^14^, and verified the results by BODIPY staining (Figure S1F). The brain LDs were 1∼2 µm in diameter and mainly localized in the cortex (Figures 1C, 1D and 1E), which is consistent with previous report^13^. In addition to the global lipid level changes, we observed the density (number), size, and distribution of LDs were all changed in aging brain (Figures S2). SRS images of the whole-mount brain show that the LDs were particularly abundant in the cell body regions of the cortex across all ages (Figures S2A). We thus quantified the LD size and density in these regions. We found that the LD density was decreased while the size was increased with aging in both sexes (Figures S2B-E). In addition, the LD density in females was consistently higher than that in the age-matched males, and the LD size in female brains was increased much more with aging than that in males (Figures 1F, 1G and S2B-E). These results demonstrate sex-dimorphism in lipid metabolism. Different from our findings of reduced LD number with aging, previous studies reported increased LD accumulation under pathological conditions, such as oxidative stress, injury, and immune challenges^10,11,13^, which suggests that lipid metabolism might be regulated by distinctive mechanisms in pathological conditions and with aging, respectively.

### DO-SRS imaging reveals reduced metabolic activities in single LDs in aged brain

LDs are dynamic organelles and heterogeneous in size and location. Previous studies show that glial LDs in *Drosophila* larval and mouse brains contain significant quantities of TAGs^13,14^. To examine whether TAGs are also the main components in LDs of *Drosophila* adult brain, we quantified LD density in *Drosophila* mutants of knockdown of glial-specific lipogenesis enzymes including Lipin and diacylglycerol acyltransferase 1 (DGAT1), and mutants of overexpression of TAG lipase (Bmm), respectively. Lipin coverts phosphatidic acids into diacylglycerols, and DGAT1 converts diacylglycerols into TAGs, while Bmm catalyzes lipolysis. We found that the brain LD densities were significantly reduced in all these 3 mutants (Figures S3F and S3G), which verified that TAGs as the main contents of LDs in adult brain.

Multi-colored SRS images at lipid-related peaks of 2850 cm^-1^, 2880 cm^-1^, and 3009 cm^-1^ showed lipids were accumulated in both cell membranes and LDs (Figures S3A), and these peaks were confirmed by SRS hyperspectral images (Figures S3B and S3C). Notably, these is no significant changes between female and male in hyperspectral shape of brain LDs and membranes, suggesting the lipid constitution is consistent during aging. Removing lipids by methanol treatment in brain tissue abolished the lipid signal at 2850 cm^-1^, and the protein signal at 2930 cm^-1^ remained (Figures S3D, S3E). Similarly, removing protein by proteinase K treatment abolished the peak at 2930 cm^-1^ (Figures S3E). These verified that the lipid signal on cell membranes was not due to bleed through from the protein signal at 2930 cm^-1^.

Remodeling of LDs is regulated by lipogenesis and lipolysis^23,24^. We thus examined the metabolic activity change in single LDs during lipogenesis and lipolysis in aging flies, respectively. We first visualized de novo lipogenesis using D_2_O probing with SRS (DO-SRS) imaging. After feeding flies at different ages (0, 10, 20, 30, 40 days posteclosion) with 20% D_2_O-containing standard diets for 5 days, we imaged LDs in the brains (Figure S4A) and quantified lipids turnover rate by the ratio of CD signal (newly-synthesized lipids) to CH signal (total lipids) collected at 2143 cm^-1^ and 2850 cm^-1^, respectively^17,18^. As shown in Figures 2B and 2C, the lipids turnover rate (CD/CH ratio) displays a bell-shaped curve in aging female and male flies. The lipids turnover gradually increased from young flies (5-15 days), reached a maximum in mid-aged flies (15-35 days), and then declined in aged flies (35-45 days). In particular, sexual dimorphism in lipids metabolism during aging was demonstrated. Lipid turnover reached a maximum of 2.4% in 25-day females but a maximum of 2.1% in 35-day males (Figures 2B and 2C). Young (5-day) males showed relatively lower CD/CH ratios than females (mean, 0.9% vs.1.4%), but old (35-day) males maintained a relatively higher rate of lipid synthesis (CD/CH ratio) compared with females (mean, 2.1% vs. 0.9%).The results show that age-related impairment of brain LD lipogenesis occurred earlier in aging females than in males. DO-SRS images also showed non-uniform distribution of newly synthesized lipids inside single LDs (Figure 2A), which is consistent with those observed in *Drosophila* fat body^18^.

**Figure 2.**
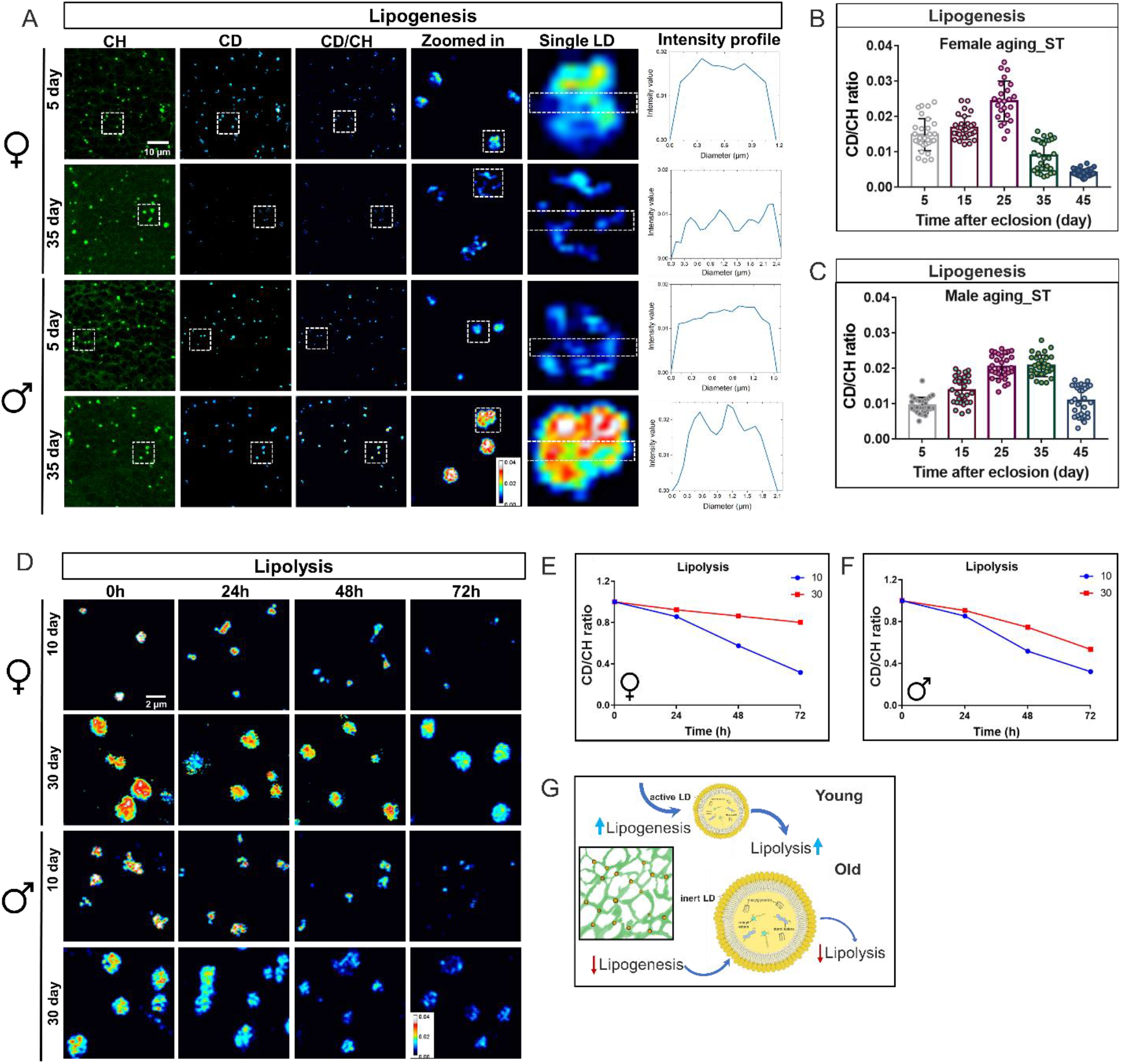
The metabolic activities of LDs in aging brains are visualized and quantified using DO-SRS imaging. (A) DO-SRS imaging of 5-day and 35-day brain LDs in female and male flies. Images were taken at 2850 cm^-1^ and 2143 cm^-1^, respectively. The signals were color-coded in green (CH, total lipids), cyan (CD, newly synthesized lipids), and royal (CD/CH), respectively. The representative LDs inside the dashed boxes were magnified. The intensity profiles inside individual LDs were plotted to show the distribution of newly synthesized lipids. Scale bar: 10µm. (B-C) Quantification of CD/CH ratios in brain LDs from female and male flies. Flies at 0, 10, 20, 30 and 40-day old were fed with 20% D2O-containing standard diet (ST) for 5 days. And then the brain LDs were imaged at 2143 cm^-1^ (CD, newly-synthesized lipids) and 2850 cm^-1^ (CH, total lipids), respectively, and the ratios of newly synthesized lipids to total lipids (CD/CH) were quantified. Error bars are Mean ± SD. n =30 ROIs from 8 brains. (D) DO-SRS images of time-course changes of lipid mobilization after 0, 24, 48, 72-hour starvation in 10- and 30-day old fly brain LDs. Scale bar: 2µm. (E-F) The quantification of CD/CH ratios of brain LDs at each time point after starvation were plotted as Mean ± SD. n =15 ROIs from 5∼8 brains. (G) A model describing the transformation of metabolic status of brain LDs during aging. The metabolically active LDs in the young brains have both high lipogenic and lipolytic rate, while the brain LDs become inert when the files grow old, which have reduced activities in both lipogenesis and lipolysis.

Brain LDs have been shown being mainly localized in glial cells^10,13^. Our DO-SRS imaging revealed strong deuterium incorporation from D2O-labeled dietary water into the core of LDs (Figure 2B and 2C), suggesting de novo fatty acid synthesis contributes neutral lipid cargo to LDs. To elucidate whether the relevant de novo lipogenesis was within the glial cells or other tissues, we used glial-specific RNAi to knock down acetyl-CoA carboxylase (ACC), the first rate-limiting enzyme for fatty acid biosynthesis. We visualized dramatic reduction of LD density in ACC knockdown flies (Figure S4C and S4D). Fatty acid synthesis is therefore required in a cell-autonomous manner for maximal induction of glial LDs. Given that induction was not completely blocked by ACC knockdown, we tested whether there was an additional contribution by hemolymph fatty acids that were directly from diets or released by peripheral tissues. Our results showed double knockdown of low density lipoprotein (LDL) receptor-related protein 1 and 2 (LRP1/2), the receptor homologues LpR1 and LpR2 responsible for uptaking hemolymph fatty acids, partially reduced the number and size of LDs (Figure S4C and 4D)^25^. Together with previous results, this indicates that both hemolymph derived fatty acids and de novo fatty acid synthesis contributed to the growth of glial LDs. Notably, the LD density was reduced to a larger extent by knockdown of ACC than LpR1/2 (Figure S4C and 4D), suggesting de novo fatty acid synthesis predominantly contributes to lipid storage under normal conditions.

As LD is a highly dynamic organelle, CD/CH values measured at each endpoint of D_2_O treatment were balanced between lipogenesis and lipolysis. To analyze the lipolysis in brain LDs, we utilized the starvation assay that facilitates the mobilization of the stored lipids. We treated 0-day (young) and 30-day old (aged) flies with 20% D_2_O for 10 days to obtain saturated deuterium incorporation (∼2.7%) for DO-SRS imaging (Figure S4B)^18^, and then transferred the flies to 1% agar for starvation. The CD/CH ratios in brain LDs were examined at 24, 48, and 72 h, respectively, after the transfer (Figures 2D-F). The CD-chasing curves indicate that lipid turnover (CD/CH) decreased faster in young flies than aged ones in both sexes (Figures 2E and 2F). LDs in 30-day old male flies had a relatively higher lipolytic rate than those in age-matched females (52% vs. 76% of initial CD/CH ratio after 72h starvation, Figures 2E and 2F). This shows that, similarly as lipogenesis, sexual difference also plays a critical role in the regulation of lipolysis during aging.

All these data indicate that, compared to young brains, LDs in aged brains have inert metabolic-activities in both lipid synthesis and mobilization (Figure 2G). The dramatic decline of lipolysis in old flies likely led to enlarged LD size, which is more significant in female brains. The general lipid loss in old brains was likely due to the reduction of lipid synthesis or misallocation of lipids in LDs. Our results demonstrated that the regulation of lipid storage in the brain is a highly dynamic process as that in the adipose tissue^18^, which is balanced between lipid synthesis and mobilization at LDs. The metabolic homeostasis resolved at the aging of single LDs might further determine the brain aging. Once this balance is disrupted, the lipid metabolic homeostasis in the brain might be disturbed, which may cause or accelerate brain aging and neuronal diseases.

### Lipogenic diets improve brain functions by promoting lipid biosynthesis

After discovering alteration of lipid metabolism in LDs of aging brain, we next investigated its association with brain functions by using the negative geotaxis climbing assay to assess the motor function. Consistent with other studies^26^, our study showed an age-dependent decline of motor activity in both females and males, and females displayed much faster decline with aging (Figures 3C and S5B). Furthermore, during normal aging process, the change of motor ability was correlated significantly with the lipogenic rate of LDs, especially at the mid (25-day) to late (45-day) age (Figure S5H).

**Figure 3.**
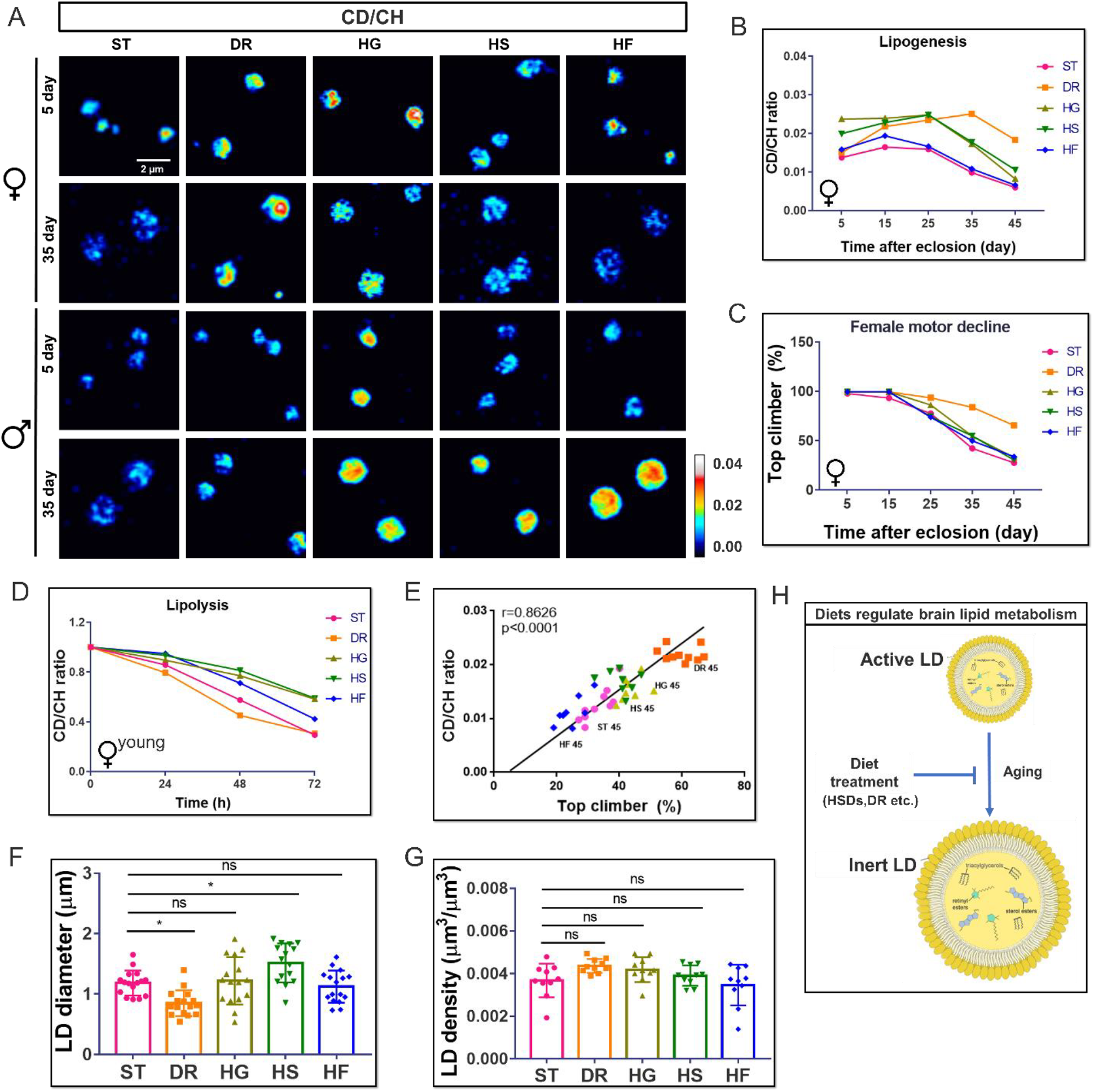
The LD metabolic activity of aging brain under diet treatments are visualized and quantified from DO-SRS imaging and it is correlated with motor activity. (A) The representative ratiometric images for diet regulated lipid biosynthesis in 5- and 35-day brain LDs (female and male) were displayed, respectively. Scale bar: 2µm. (B) The brain LDs from 5, 15, 25, 35 and 45-day old females pre-treated with 20% D2O labeled standard diet (ST), high sugar (glucose, fructose and sucrose) diets (HSDs) and dietary restriction (DR) for 5 days were imaged from the CD lipids (2143 cm^-1^), CH lipids (2850 cm^-1^), and the ratio of newly synthesized lipids to total lipids (CD/CH) during 5-day D_2_O-labeling were quantified and plotted as Mean ± SD. n =15 ROIs from 5∼8 brains. (C) The motor activities of 5, 15, 25, 35 and 45-day old females fed with standard diet (ST), high sugar (3xglucose, fructose and sucrose) diets (HSDs) were evaluated and plotted as Mean ± SD. N = 40 flies for each age group. (D) The time-course changes of CD-lipid mobilization over 72-hour starvation in LDs from 10-day young females were quantified and plotted as Mean ± SD. n =15 ROIs from 5∼8 brains. (E) A linear positive correlation between brain lipid synthesis and motor ability of 45-day old flies under different diet treatment. n = 10∼15 for each group, Pearson analysis, r = 0.8626, p < 0.0001. (F) and (G) Quantitative comparison of LD density (F) and diameter (G) in diet manipulated 35-day fly brains. Results were plotted as Mean ± SD and compared. N =8∼10 brains for each age group. Statistical significance was determined by 2-way ANOVA. *, p < 0.05; **, p < 0.01; ***, p < 0.001; ****, p < 0.0001; ns, non-significant difference.

Although the molecular pathways regulating brain lipid homeostasis and concomitant motor deficits are unclear, dietary intervention has been considered an efficient approach to promote motor ability and extend healthy lifespan^27-30^. We first observed flies treated with high sugar diets (HSDs, 3x glucose, sucrose, or fructose) or dietary restriction (DR, 0.5x yeast) had notably extended lifespans in both sexes (Figures S5C, S5D). We further demonstrated that the motor activities were affected at various degrees under diet treatments (Figures 3C and S5B). DR can significantly increase the later-stage mobility (25∼45-day) in both sexes, while HSDs benefit males more (Figures 3C and S5B).

We then studied the correlation between diet regulated motor function and brain lipid metabolic activity. We examined lipid metabolic changes in fly brains under various diet treatments, including standard food (ST), dietary restriction (DR), high glucose (HG), high sucrose (HS) and high fructose (HF). The lipogenesis of brain LDs were systemically imaged (Figures 3A and S6) and quantitated in female (Figure 3B) and male (Figure S5A) flies on different diets and at different ages. The lipid synthetic rate increased first and then declined in aged flies under all diet treatments. However, HG and HS diets, as well as DR, greatly enhanced lipogenesis in female flies at all ages, and HF diet only slightly increased lipid biosynthesis in female files (Figure 3B and S6). All these three sugar diets significantly enhanced lipogenesis in male flies (Figure S5A). The lipogenic effects of different sugar diets on brains (glucose/sucrose > fructose) are different from those on fat bodies, in which HS and HF diets exerted greater effects at different ages than HG diet (fructose /sucrose > glucose) (Figure S7). This suggests that different types of sugar might be sensed and processed by different pathways in specific tissues.

Among all diet treatments, DR led to the most significant enhancements of lipogenesis and motor ability in both old female and male brains (45-day). Particularly, lipogenesis increased more in DR treated 45-day old females than males (3.6-fold vs. 2.1-fold), suggesting DR can promote more lipids synthesis in females. This is consistent with the lifespan data (Figures S5C and S5D) that DR significantly extended the median lifespan of females by 28% but had no effects on males. High sugar diets had particularly marked impacts on male’s lifespan, with a 32% extension under high glucose treatment (Figures S5C and S5D). However, DR still had the most impact on lipogenesis in 45-day old males. Furthermore, the lipogenesis in aged brain (45-day) correlated significantly with motor ability (Figure 3E) under all diet treatments.

To examine how lipid mobilization in brain is altered by diets, we performed starvation assays for 72 h on 10-day (young) and 30-day (old) female and male flies pretreated with HSDs or DR, respectively. The lipolytic curve showed that young flies mobilized lipid storage faster than old ones, and males faster than females under all diet pretreatments (Figures 3D and S5E-G). In both sexes, young flies pretreated with HSDs had lower lipid mobilization rate than control, suggesting the metabolic pathways might be reprogrammed under HSD treatments. In young files, HF led to a relatively higher lipolytic rates, but HG and HS diets showed no significant differences. This again underscored that metabolism affected by fructose was differently from the other two sugar diets. In old flies, HSDs-pretreatment showed no significant differences in lipolysis compared with normal ones (Figures S5F and S5G), indicating the signaling activity of metabolic pathways involved in HSDs regulation may be altered during aging. Nevertheless, both young and old flies pretreated with DR mobilized lipids rapidly during 72 h starvation, and was the most significant in old females with an endpoint change of 0.46-fold to that of 0.81-fold in control *w*^*1118*^ females (Figure S5F and S5G).

Taken together, diets regulated brain lipid metabolism by modulating lipogenesis and lipolysis in LDs (Figure 3H), without significant changes in LD size and density (Figures 3F and 3G), which might be closely correlated with brain functions and lifespan. Our HSDs and DR recipes contained relatively large amounts of sugar (sugar to protein ratio 3:1 and 1:1, respectively), compared to the standard diet (sugar to protein ratio 0.5:1) (Table S1). They exerted similar lipogenic effects on brain metabolism in old flies, which may benefit the flies’ lifespan by preventing lipid loss as brain aging. This is supported by the study that dietary balance of protein to carbohydrate (P:C) ratio, instead of the total calorie, is the predominant determinant for the lifespan^30^. In addition to lipogenesis, DR pretreatment also significantly increased the lipolytic rate under starvation, suggesting the basal lipid metabolic pathways may be readapted to promote motor activity and lifespan to the maximum extent.

### IIS pathway mediates brain lipid metabolism

The insulin/IGF-1 signaling (IIS) pathway is a primary regulator of development and aging by controlling metabolic homeostasis of carbohydrates and lipids^31-34^. Dysregulation of the IIS pathway can lead to insulin resistance, diabetes, and aspects of metabolic disorders^35, 36^. Reduced IIS activity can extend lifespan in worms, flies and mammals (Figure 4A)^37^, as well as improve the stress resistance to environments^31, 38^.

**Figure 4.**
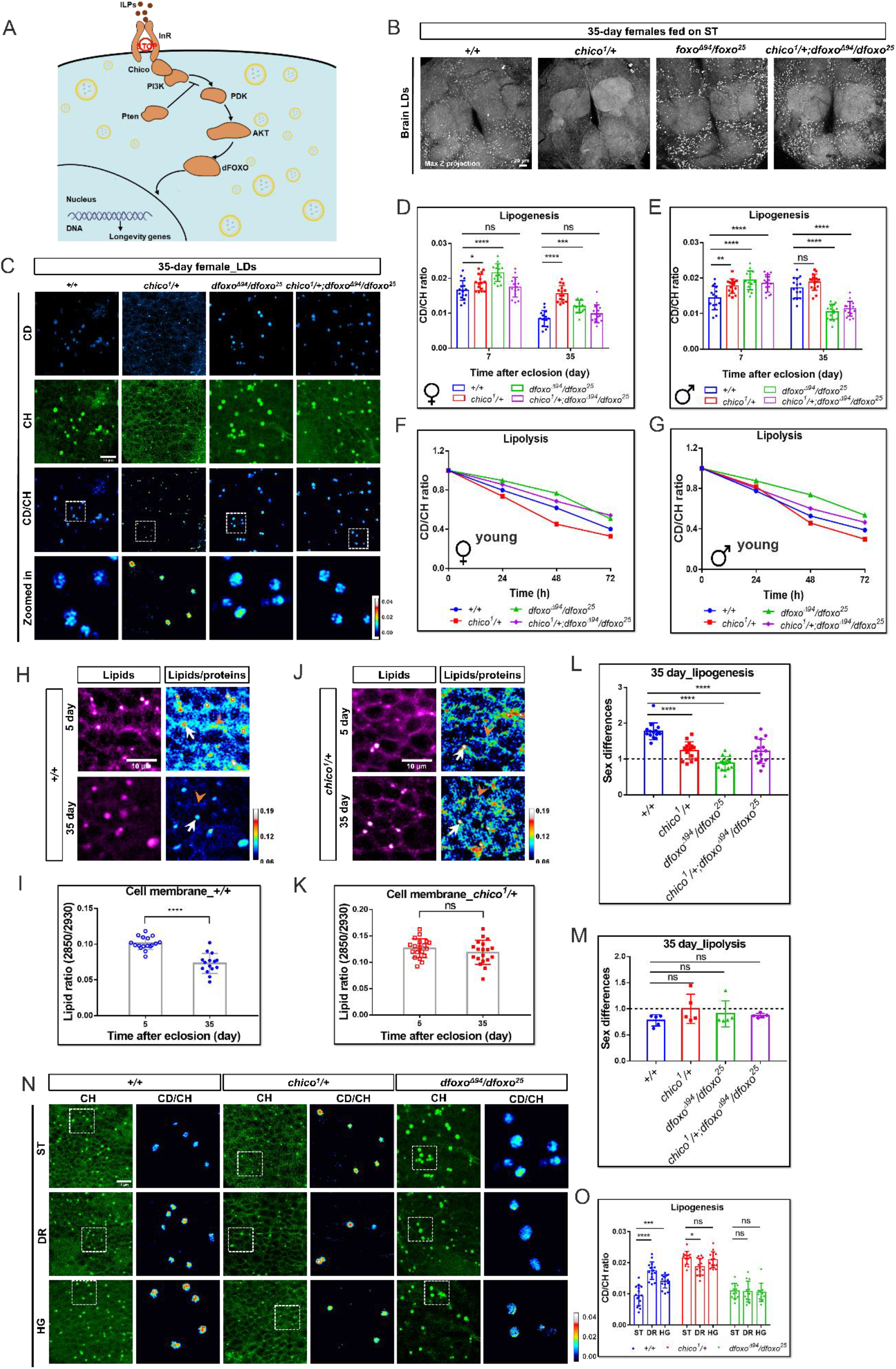
The lipid metabolic changes in brains of IIS mutant flies. (A) A schematic model depicting the key components of IIS pathway regulating longevity in *Drosophila*. When InR is inactive or Pten is active, the signaling activity is inhibited, the dephosphorylated dFOXO will be translocated to the nucleus to active the expression of downstream longevity related genes. (B) Max Z projected images of brain LDs from IIS mutants (*chico1/+, dfoxo*^*Δ94*^*/dfoxo*^*25*^and *chico1/+; dfoxo*^*Δ94*^*/dfoxo*^*25*^) and control (*w*^*1118*^). (C) DO-SRS images of brain LDs from 35-day old IIS mutant flies. Files were fed with 20% D2O labeled standard diet (ST) for 5 days. Images from the CD lipids (newly synthesized lipids, 2143 cm^-1^) and CH lipids (total lipids, 2850 cm^-1^) were displayed. Ratiometric images (CD/CH) show the lipid turnover rate. The representative LDs inside the dashed box were magnified to show the details. Scale bar: 10µm. (D-E) Quantification of lipid biosynthesis in brain LDs from 7- (young) and 35-day (old) female (D) and male (E) mutants. n =15 ROIs from 3∼5 brains. (F-G) The time-course quantification of CD-lipid mobilization (CD ratio decline) in 10-day old (young) female (F) and male (G) brain LDs after 0, 24, 48, 72-hour starvation were plotted as Mean ± SD. n =15 ROIs from 3∼5 brains at each group. (H-K) The SRS images of 2850 cm^-1^ as shown in magenta hot were to show distribution of lipids in subcellular level of cell body regions from 5- and 35-day old control flies (*+/+*) (H) and IIS mutants (*chico*^*1*^*/+*) (J). The ratiometric images of 2850 cm^-1^ /2930 cm^-1^ was shown as royal to quantitatively mapping the lipids level in subcellular level of cell body regions. White arrows, LDs; orange arrow heads, cell membranes. Scale bar: 10 µm. The quantitative results were plotted as Mean ± SD in (I) and (K). (L-M) The sex differences of lipid metabolism in IIS mutants were determined. The sex differences in lipogenesis (L) and lipolysis (M) (arbitrarily defined as endpoint of M_CD/CH_/F_CD/CH_) were quantified and compared between control and IIS mutant flies. (N) The representative SRS images at CH lipids (2850 cm^-1^), and ratiometric images for lipid biosynthesis (CD/CH ratios) in control fly, *chico*^*1*^*/+* and *dfoxo*^*Δ94*^*/dfoxo*^*25*^ mutant brain LDs under diet treatments (ST, DR, HG) were displayed. Scale bar: 10 µm. (O) Quantification of lipid turnover in *chico*^*1*^*/+, dfoxo*^*Δ94*^*/dfoxo*^*25*^, and control flies under HG, DR, and standard diet treatments. Statistical significance was determined by using 1-or 2-way ANOVA. *, p < 0.05; **, p < 0.01; ***, p < 0.001; ****, p < 0.0001; ns, non-significant difference.

To further understand the role of IIS in brain lipid metabolism during aging, we studied *Drosophila* mutants with reduced functional copies of *chico*, a homolog of insulin receptor substrate (IRS) that mediates IIS. The heterozygous adults with a loss of function allele, genotypically *chico*^*1*^*/+*, were shown to have extended lifespan and promoted motor activities^39-42^. We found that the density and size of brain LDs were reduced significantly in both female (Figures 4A, S8A and S8B) and male *chico*^*1*^*/+* flies (data not shown). This reduction reflects a decrease of the average lipid storage in brain, which could either be due to less fat accumulation or increased lipolytic activity.

To elucidate this, we examined both the lipogenesis and lipolysis in *chico*^*1*^*/+* mutant fly brain. Interestingly, SRS images and quantification results indicate that, compared to *w*^*1118*^ (*+/+*) female controls, the lipogenic rates in LDs increased significantly in both 7-day (by 0.15-fold) and 35-day (by 0.85-fold) female *chico*^*1*^*/+* fly brains (Figures 4C and 4D). Similar effects were observed in 7-day and 35-day male flies, with an increase of 0.24-fold and 0.09-fold, respectively (Figure 4E, image not shown). These results show that LDs in *chico*^*1*^*/+* fly brain maintained high lipogenic activity during aging. By starvation tests on flies for 72 h, we found the lipid turnover rates (CD/CH ratios) were significantly decreased in both young and old *chico*^*1*^*/+* flies (Figures 4F, 4G, S8D and S8E), indicating enhanced lipid mobilization. Together, we concluded that both lipogenesis and lipolysis activities increased in *chico*^*1*^*/+* flies, and the dramatically reduced density and size of LD in *chico*^*1*^*/+* brain were due to much higher lipolytic activities than lipogenesis. Moreover, we found that lipid contents in brain membrane were also significantly reduced with aging in control flies (Figures 4H and 4I), but were maintained in IIS downregulation (*chico*^*1*^*/+*) flies (Figures 4J and 4K). Hypothetically, the reduced IIS activity might help maintain brain function in youth-status during aging by facilitating membrane lipids turnover.

Downregulated IIS activity has been shown to improve metabolic profile in *Drosophila* fat body and muscle, mainly by working through the downstream transcription factor dFOXO (Figure 4A)^31, 43-48^. We then examined how lipids metabolism was modulated by dFOXO in the brain, by using *dfoxo*^*Δ94*^ and *dfoxo*^*25*^ *Drosophila* mutants^49^ (*dfoxo* loss of function). Contrary to those smaller LDs observed in *chico*^*1*^*/+* flies, LDs in *dfoxo* mutants were much larger than control (Figures 4B, 4C and S8B), which could be due to increased lipid biosynthesis, reduced lipolysis, or both. Our quantification results of lipogenesis indicated that, compared to the control, the lipogenic rates were increased by 0.31-fold and 0.42-fold in 7-day and 35-day female *dfoxo*^*Δ94*^/*dfoxo*^*25*^ mutants, respectively, and 0.33-fold in 7-day male *dfoxo* mutants (Figures 4D and 4E). However, in 35-day male *dfoxo* mutants the lipogenic rate was significantly decreased by 0.32-fold (Figure 4E). These results indicate that female and male flies have different susceptibility to lipid metabolic defects caused by *dfoxo* loss of function. More interestingly, the sex differences (the ratio of CD/CH in male to that in female with respective genotype) of lipogenesis in 35-day IIS mutants (*chico*^*1*^*/+* and *dfoxo*) were significantly reduced (approximate to 1) compared with control (Figures 4L and S8C).

We next examined the lipolytic rates in these *dfoxo* mutants with starvation assays for 72h. The CD/CH ratios in *dfoxo* mutants were declined during 72h starvation in both female and male flies, but not as much as those in the control (Figures 4F, 4G, S8D and S8E). These data together indicates that the metabolic activity in old *dfoxo*^*Δ94*^/*dfoxo*^*25*^ mutants’ brain was reduced and the enlarged LDs in the brain were due to less lipolysis compared with control. The sex differences of lipolysis were slightly but not significantly reduced (approximate to 1) in aged *chico*^*1*^ and *dfoxo* mutants compared to the control (Figure 4M). We also compared the LD size and lipogenic activities in 0-day (the day flies eclosed from pupae) *dfoxo*^*Δ94*^/*dfoxo*^*25*^ mutant with control and found no significant differences, suggesting the change of lipid metabolism was not from development but adult-onset (Figure S8F and S8G). Altogether, these data indicate that IIS pathway may be involved in the regulation of sex differences in brain lipid metabolism.

We further conducted metabolic assays on combined *chico*^*1*^*/+; dfoxo*^*Δ94*^*/dfoxo*^*25*^ mutant flies. DO-SRS imaging and quantification of lipogenesis show similar results as those in *dfoxo* mutants (Figures 4A-E, S8A and S8B), indicating IIS pathway functions through dFOXO to regulate brain lipid metabolism.

Study showed that IIS is low when nutrients are sparse and dFOXO regulates a positive feedback loop to increase *Drosophila* insulin receptor (dInR) at the cell surface^50^. Therefore, the sensitivity of the InR signaling pathway could be increased by dFOXO activity, which probably results in the increased lipogenic rate as in our study. On the other hand, lipases such as Bmm are the metabolic targets of dFOXO, so when dFOXO is translocated to the nucleus and IIS is downregulated, the transcription of the lipase is active, and the lipolysis rate is increased. This is likely the mechanism that both the lipogenic and lipolytic rate were increased as we observed in DR and *chico*^*1*^*/+* flies.

Previous studies showed that removing one functional copy of *chico* or loss function of *dfoxo* did not extend the lifespans of flies on high sugar diets^26, 51^. We examined the effects of diet treatments on brain metabolism in IIS mutants. We found that *chico*^*1*^*/+* flies fed on HG or DR had no significant difference in lipogenesis and LD size, compared with those mutants on standard diet (Figures 4N and 4O). In addition, the metabolic activity of enlarged LDs in *dfoxo* mutant flies could not be further modulated by HG and DR (Figures 4N and 4O). All these results together demonstrate that IIS-dFOXO axis plays a crucial role in mediating diet-regulated lipid metabolism of *Drosophila* brain.

### Downregulation of IIS in glial cells promotes lipid turnover

Glia-specific IIS inhibition was shown to extend lifespan in *Drosophila*^52^. *To investigate how local brain lipid metabolic activity was regulated by the IIS in adult brain (not in the developmental brain). We applied mifepristone (RU486)-inducible glia specific Gal4 driver GSG907* to reduce IIS activity in adult brain, including *GliaGS>UAS-Pten* (a lipid phosphatase counteracting PI3K enzymatic activity), *GliaGS>UAS-InRDN* (the dominant negative form of InR), *GliaGS>UAS-InRCA* (constitutively active form of InR) flies, and *GliaGS>UAS-lacZ* as control. Both *GliaGS>UAS-Pten* and *GliaGS>UAS-InRDN* mutants showed reduced LD density phenotype in brains (Figures 5A and 5B), recapitulating that observed in *chico*^*1*^*/+* flies (Figures 4B, S8A and S8B), even though to a lesser extent.

**Figure 5.**
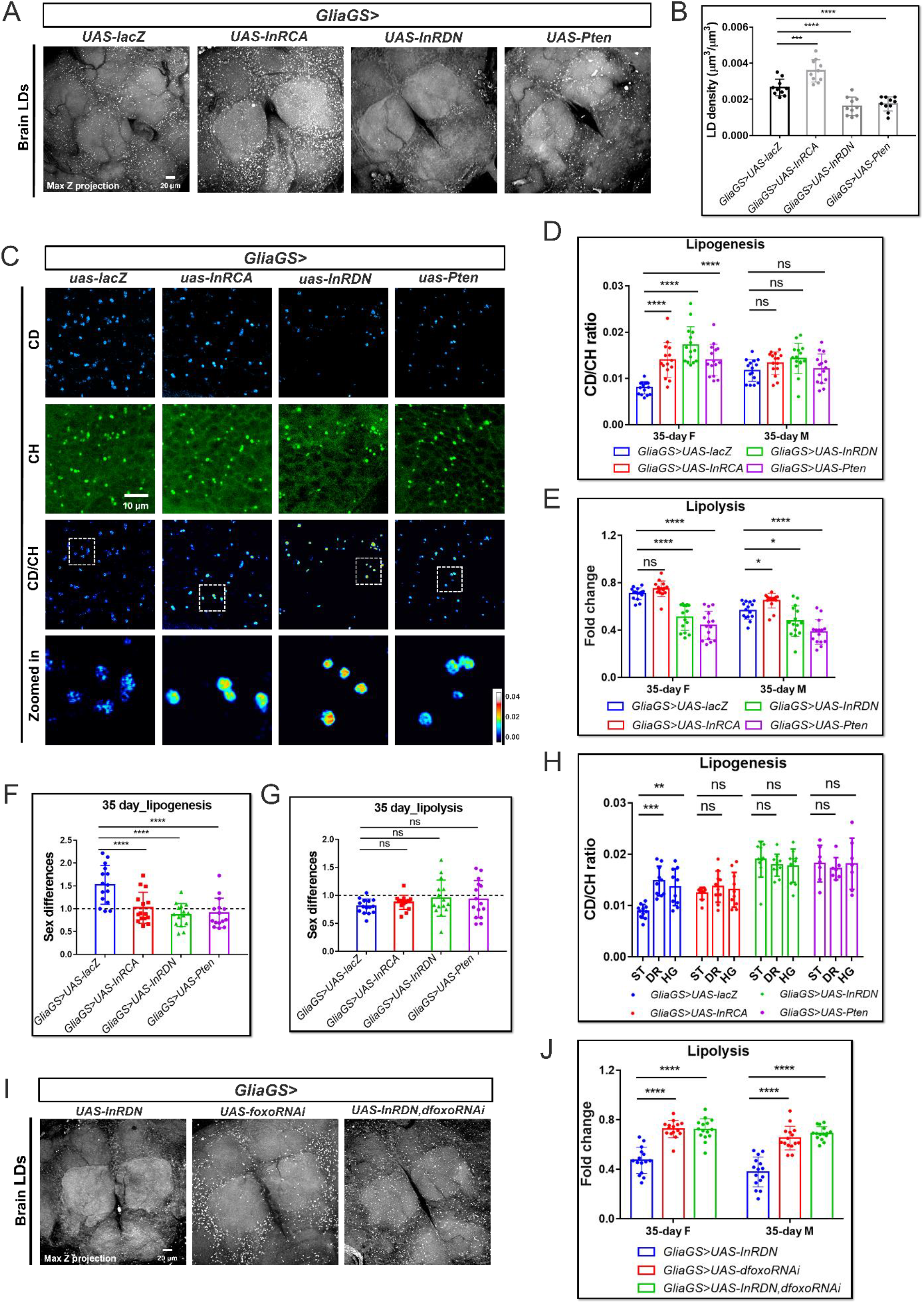
Glia-specific manipulation of IIS pathway modulates brain lipid metabolism. (A) Max Z projected images of brain LDs from glia-specific gene manipulated (*GliaGS>*) adult flies (*UAS-InRCA, UAS-InRDN* and *UAS-Pten*) are compared with that of *GliaGS>UAS-lacZ*. (B) The LD density ratios (total LD volume to brain cortex volume) in (A) were quantified and presented as Mean ± SD. n = 5 brains in each group. (C) DO-SRS images of LDs in 35-day IIS mutant flies. The representative LDs inside the dashed box were magnified to show the details, Scale bar: 10µm. The lipid turnover rates (CD/CH) were quantified and plotted as Mean ± SD in (D). n =15 ROIs from 3∼5 brains in each group. (D) Quantification of lipid turnover in 35-day female and male mutants. All female mutants showed significantly higher lipogenesis compared with control, while there were no significant changes in male mutants. (E) The quantification of CD-lipid mobilization as endpoint fold changes from IIS genetic manipulated (genotypes as shown in the figures) 35-day old female and male brain LDs after 72-hour starvation were plotted as Mean ± SD. n =15 ROIs from 3∼5 brains in each group. (F-G) The sex differences of lipid metabolism in glia specific IIS manipulated flies were determined. The sex differences in lipogenesis (F) and lipolysis (G) (arbitrarily defined as endpoint of M_CD/CH_/F_CD/CH_) were quantified and compared between control and glia specific IIS manipulated flies. (H) Quantification of lipid turnover in glia specific IIS manipulated flies and control flies under HG, DR, and standard diet treatments. (I) Max Z projected images of brain LDs from glia-specific gene manipulated (*GliaGS>*) adult flies (*UAS-dfoxoRNAi* and *UAS-InRDN,UAS-dfoxoRNAi*) were compared with that of control (*GliaGS>UAS-lacZ*). n = 3∼5 brains in each group. (J) The quantification of CD-lipid mobilization as endpoint fold changes in brain LDs from 35-day old IIS mutants after 72-hour starvation. Data were plotted as Mean ± SD. n =15 ROIs from 3∼5 brains in each group. Statistical significance was determined by using 1-or 2-way ANOVA. *, p < 0.05; **, p < 0.01; ***, p < 0.001; ****, p < 0.0001; ns, non-significant difference.

As the young flies fed with RU-486 takes time to induce the expression of GAL4 and the downstream targets and the effects from genetic manipulation is not stable. Therefore, we examined brain lipid metabolic activities in 35-day (after a long time induction of target gene expression) mutants, including *GliaGS>UAS-Pten, GliaGS>UAS-InRDN, GliaGS>UAS-InRCA* flies, and *GliaGS>UAS-lacZ* as control. DO-SRS ratiometric images showed much more CD/CH signals in the LDs of 35-day old *GliaGS>UAS-Pten, GliaGS>UAS-InRDN*, and *GliaGS>UAS-InRCA* flies, compared with *GliaGS>UAS-lacZ* control flies (Figure 5C). Quantification results of lipogenesis showed that downregulation of IIS downstream component (*GliaGS>UAS-Pten*) or upstream receptor (*GliaGS>UAS-InRDN*) increased lipid biosynthesis significantly but the LD density was decreased compared with control (*GliaGS>UAS-lacZ*), while constitutively active IIS (*GliaGS>UAS-InRCA*) increased both lipid biosynthesis and LD density, which is probably because its promotion of anabolic processes (Figures 5A-D). Surprisingly, there were no significant changes of lipogenesis in all male mutant flies (Figure 5D), showing that females are more sensitive to genetic manipulations of IIS.

Further comparison of lipid catabolic rate in *GliaGS>UAS-InRCA* aging flies revealed decreased ability to mobilize lipid storage (higher CD/CH ratio) compared with the control; while genetic downregulation of IIS pathway (*GliaGS>UAS-Pten* and *GliaGS>UAS-InRDN*) significantly decreased CD/CH ratios, correspondingly, higher lipolytic rate in the old flies (Figure 5E), which shows the same trend as in old *chico*^*1*^*/+* flies (Figures 4F and 4G). Consistent with the systematic downregulation of IIS, glia-specific IIS downregulation reduced the sex differences significantly in lipogenesis, but no significant changes in lipolysis (Figures 5F and 5G). The DR- and HG diet-induced lipogenesis enhancements were abolished in all mutant flies (Figures 5H).

To test if the activity of dFOXO was upregulated upon reduced IIS in glial, which could facilitate the lipolysis, we examined the lipid metabolic dynamics in *dfoxo*RNAi crossed *GliaGS>UAS-InRDN* flies (*GliaGS>UAS-InRDN,UAS-dfoxoRNAi*), as well as *GliaGS>UAS-InRDN* and *GliaGS>UAS-dfoxoRNAi* flies, respectively, for comparison. We found significantly higher LD density in *GliaGS>UAS-dfoxoRNAi* and *GliaGS>UAS-InRDN,UAS-dfoxoRNAi* flies, which had greatly reduced lipolysis (higher CD/CH ratio), compared with *GliaGS>UAS-InRDN* control (Figures 5I and 5J). But there was no difference between these two mutants, showing dFOXO was is epistatic to InRDN overexpression. These results indicate dFOXO is responsible for IIS in glial cells to regulate brain lipid metabolism.

We further verified the effects of IIS downregulation on brain lipid metabolism were glial specific. Using the neuron-specific *elavGS* driver to downregulate IIS in mutant flies (*elavGS>UAS-InRDN*), we found there were no significant changes of LD density and lipogenesis compared with control (Figures S9A-C). Altogether, we showed that glia-specific IIS autonomously regulates the lipid metabolism of *Drosophila* aging brain.

## Discussion

For the first time we employed DO-SRS imaging to directly visualize and quantitate the spatiotemporal changes of lipid metabolic activity in aging *Drosophila* brain and examined the interconnections between brain lipid metabolism, dietary composition, IIS pathway, and motor activity. Recent studies have shown that lipid metabolism in brain not only is associated with aging but also actively participates in the regulation of aging and lifespan^2, 11, 12, 14, 20, 53-60^. Diet interventions can modify the locomotor activity and healthy lifespan in model systems^26, 27, 29, 61-63^, but the underlying mechanisms remain elusive. Our study identified that glia IIS pathway mediated brain lipids metabolic turnover, which bridges this critical knowledge gap.

In normal physiological situations, there are numerous metabolically active LDs in young brains. They probably need high activity on biosynthesis and lipolysis to readily provide lipid materials and energy sources to neuronal and glial cells. It has been reported that lipid shuttling between neuron and glia contributes to both the development and function of the neuron, as well as lipid homeostasis in the brain^57, 58", 64-70^. Previous studies showed fatty acids from membrane phospholipids could redistribute to LDs during oxidative stress^13, 71^, suggesting a mutual interaction may exist between LDs and membranes. Moreover, we detected reduced lipids level in cell membranes combined with accumulated lipids in enlarged LDs during aging (Figures S3A, 2B and 4J), which is likely due to impaired lipid exchanges between LD storage and membranes in aged brain (Figure 6). This might be one of the major reasons that causing brain dysfunctions^56, 72^. Furthermore, the rates of lipid biosynthesis and mobilization were decreased in the inert LDs of aged brains. It probably also affects the efficiency of lipid shuttling between neuron and glia, which causes the energy shortage in the neuronal system.

Our results consistently show sexual dimorphism in regulating brain lipid metabolism during aging^62, 73, 74^. In general, males maintain a higher lipid metabolic rate (both in lipogenesis and lipolysis) than females, especially during old stages, which may be correlated with their higher motor activity. We found that DR treatment improved lipid metabolic activity in female brains, while HSDs were more beneficial to male brains. These sex-dependent effects are more likely mediated by the IIS pathway, as shown by the sexual differences of LD dynamics being abolished in *chico*^*1*^*/+* and downregulated IIS mutant flies (Figures 4H, 4J, S8C, 5F and 7G). Previous studies have found IIS plays a sex-dependent role in the body size and locomotor activity, as well as susceptibility to develop diabetes^40, 75-79^. We further uncovered that dFOXO-GFP shows different expression patterns between female and male fat bodies, with more nuclear localized dFOXO-GFP in females (Figure S10A). However, the spatial distribution of dFOXO-GFP in female and male brains showed no significant difference. Nevertheless, reduced IIS can affect aging by modulating dFOXO activity independently of subcellular localization. It is also possible that, IIS-dFOXO axis may not be the most dominated payer in regulating the sex difference in brain metabolism, but some other systematic or local signaling pathways participate in this process. For example, Ecdysone-EcR pathway, which has been shown to regulate lipid metabolism in female *Drosophila* ovary^80^, may also function in brain lipid metabolism in a sex-dependent manner. Therefore, the complex mechanism underlying the sexually dimorphic regulation of brain lipid metabolism under diet treatments needs further investigation.

Continuously high levels of IIS over the life span would lead to deleterious consequences and contribute to pathological changes associated with age through the effect on cell division and/or metabolism^81^. In fact, although the lipid loss was prevented and the brain function was improved by lipid synthesis increase under HSDs, the activity of lipid mobilization was decrease, which might be detrimental to the organisms and the main reason causing later-stage death^35, 63, 82^. Ectopic activation of IIS through overexpressed InRCA in glia phenocopies the influences induced by HSDs, causing increased lipogenesis and decreased lipolysis, as well as brain lipid accumulation. These results together agree with the recent theory on the evolution of aging that the expression of particular genes benefits biogenesis early in life but will become detrimental with age^83^. On the other hand, inhibition or downregulation of IIS in glia promoted lipid metabolic activity in *Drosophila* brain, which plays an important role in extending the lifespan^52, 84^. It also underscores the crucial role of brain lipid metabolism in maintaining not only nervous system function but also organism-wide health. It has been shown that dFOXO mediates reduction of IIS in different tissues to regulate *Drosophila* healthy lifespan^43, 44, 46, 85^. Our study found that overexpression of InRDN in glia cells improved lipid metabolic activity, which was reversed by *dfoxo* knockdown. Glia-specific knockdown of *dfoxo* alone is sufficient to induce large number of LDs by reducing the lipolysis. Moreover, glia-specific manipulation of IIS pathway causes metabolic insensitivity to nutritional regulation^33, 86^. The regulatory function of dFOXO in *Drosophila* brain lipid metabolism is consistent with recent study in mouse model of amyloid pathology^87^, suggesting the interconnections between IIS, glia lipid homeostasis and Alzheimer’s diseases.

In summary, our study presents a new approach of optical imaging to directly visualize and quantify spatiotemporal alterations of lipids turnover in *Drosophila* brain, providing a better understanding of the impacts of IIS pathway on brain lipid metabolism, and showing high metabolic activities in the brains are correlated with the motor abilities and healthy lifespan. Our study also provides important insights into the sexual differences of brain lipids metabolism, which might be modulated by IIS activity.

## Limitations of the study

Our study found that diets regulate brain lipid metabolism by modulating glia IIS/dFOXO pathway. The newly synthesized fatty acids are well tracked by D_2_O labeling under different nutritional and genetic conditions. However, we did not trace how lipids were utilized in these different contexts. Under the situations of prolonged starvation or reduced IIS activity, the lipid catabolism was dominated, suggesting lipogenesis and lipolysis were readjusted and different from that in steady state. Whether the reprogrammed metabolism itself or the downstream substances from lipolysis, such as ketones^88^, can help neuron survival, thereby promoting the lifespan is an important issue that needs to be further explored.

We hypothesized a mutual communication exists between membrane lipids and LDs and it is enhanced by downregulation of IIS activity. However, due to technical obstacles, measuring the dynamic change of lipid membrane and LDs is a major challenge. In addition, the exact mechanism that regulates the membrane-LD translocation is still unclear. Genetic screening can be used for mechanistically precisely characterizing this IIS-related regulatory mechanism.

Our data suggested a crucial role of pan-glial cells playing in brain lipid homeostasis, but the function of different subtypes of glia and neuron in regulating brain lipid metabolism and healthy lifespan is unknown. It is likely that different subtype of glial/neuronal cells has distinct metabolic activity and molecular profiles. The label-free in situ metabolic imaging can be used to map the brain metabolic atlas, which can be a reference to the human brain study.

## Acknowledgements

We are grateful to Bloomington Drosophila Stock Center for fly stocks. We thank Drs. K. Zhang, C. Metallo, G. Haddad, D. Zhou, and Shi Lab group members for helpful discussions. We acknowledge support from UCSD Startup funds, NIH U54 2U54CA132378, NIH 5R01NS111039, NIH R21NS125395, NIHU54DK134301, NIHU54 HL165443, and Hellman Fellow Award.

## Author contributions

L.S. and Y. L. conceived the idea, designed the study, interpreted data, and wrote the paper. Y.L. conducted the experiments, analyzed the data, and performed statistical analyses with the help from P.C., S.S., H.J., Y.N., A.Z., S.H., X.C. and L.S.

## Materials and Methods

### Drosophila genetics

Fly lines used in this study were originally obtained from the Bloomington Drosophila Stock Center (BDSC) unless otherwise stated. They have been maintained on standard diet (Nutri-Fly, Cat #: 66-113, Genesee Scientific Corporation) in the lab for several generations. The wild type used for normal aging and diet treatment was *w*^*1118*^(stock #5905). Genetic elements used were *repo-Gal4* (stock #7415), *GliaGS* (*GSG550*, stock #62085 or *GSG907*, stock #40310), *elavGS* (kindly provided by Dr. Xu Chen), *UAS-lacZ* (stock #8530), *UAS-InRCA*(also known as *InRdel*, stock #8248), *UAS-InR*.*DN* (stock #8253), *UAS-Pten* (stock #82170), *UAS-ACC RNAi* (stock #34885), *UAS-white RNAi* (was crossed out from stock #65409), *UAS-LpR1 RNAi* (stock #50737), *UAS-LpR2 RNAi* (stock #54461), *UAS-Lipin* RNAi (kindly provided by Dr. Jane Wu, stock #63614), *UAS-DGAT1RNAi* (stock #65963), *UAS-Bmm* (stock #76600), *UAS-dfoxo RNAi* (stock #32427). IIS pathway mutant alleles and reporter used were *chico*^*1*^ (stock #10738), *dfoxo*^*25*^(stock #80944), *dfoxo*^*Δ94*^(stock #42220) and dFOXO-GFP (stock #38644).

### Stimulated Raman Scattering Microscopy

An upright laser-scanning microscope (DIY multiphoton, Olympus) with a 25x water objective (XLPLN, WMP2, 1.05 NA, Olympus) was applied for near-IR throughput. Synchronized pulsed pump beam (tunable 720–990 nm wavelength, 5–6 ps pulse width, and 80 MHz repetition rate) and Stokes (wavelength at 1032nm, 6 ps pulse width, and 80 MHz repetition rate) were supplied by a picoEmerald system (Applied Physics & Electronics) and coupled into the microscope. The pump and Stokes beams were collected in transmission by a high NA oil condenser (1.4 NA). A high O.D. shortpass filter (950 nm, Thorlabs) was used that would completely block the Stokes beam and transmit the pump beam only onto a Si photodiode for detecting the stimulated Raman loss signal. The output current from the photodiode was terminated, filtered, and demodulated by a lock-in amplifier at 20 MHz.The demodulated signal was fed into the FV3000 software module FV-OSR (Olympus) to form image during laser scanning. All images obtained were 512 × 512 pixels, with a dwell time 80 μs and imaging speed of ∼23 s per image. A background image was acquired at 2190 cm^-1^ and subtracted from all SRS images using ImageJ.

### LD analysis

3D *Drosophila* brain images were taken by SRS imaging system at 2850 cm^-1^. To enhance the detection precision, the pixel numbers were increased by 2 times along the lateral direction and 4 times along the axial direction and then A-PoD ^89^ was used to convert the preprocessed 3D images to the super-resolved images. From the super-resolved image, the numbers and sizes of LDs were measured with the 3D objects counter plugin ^90^ of ImageJ. The information about volume and location of each LD were exported and delivered to a home-built Matlab code to visualize the data as color-coded 3D images. To reduce the background noise, band pass filter (threshold: 50 px) was applied to filtering high-frequency signal. The profile of the central brains including neuropiles were delineated by FeatureJ structure plugin ^91^. The color-coded LD images and central brain areas are combined into a single image stack to present the 3D distribution of each LD in the *Drosophila* brain.

### Climbing Assays

The motor neuron functions of flies were determined by climbing assays as reported^52^. Parental flies (*w*^*1118*^) were kept for 15 females and 10 males per vial and flipped every 2 days to prevent overcrowding. Progeny flies were collected within 24 h of eclosion and aged without further CO2 exposure. After 2 days’ maturation, females and males were separated to different groups. For each diet condition, 6 vials of each sex with 20 flies per vial (totally 120 females, 120 males) were set up for examining the changes of motor activities during aging process. Flies were flipped onto fresh food every 2∼3 days. Each data point represents 15∼20 flies (natural death happens in each vial during aging) in one vial. Climbing experiments were carried out approximately at the same time during the daytime to minimize circadian differences. Flies were transferred to the glass cylindrical vials with a line at the 5 cm mark and allowed to acclimate the circumstance for 10∼15 min before testing. Flies were lightly tapped down to the bottom of the vial, and the number that crossed the 5 cm mark in 30 s were counted. Vials were placed horizontally and retested 10∼15 min later. The average of the two technical replicates for each vial was recorded, and the percentage climbing was plotted as a single point. The climbing analysis were carried out at 5, 15, 25, 35, 45 days after eclosion.

Comparisons were made using either independent samples t-tests or, in the case of three or more means, one-way analysis of variance (ANOVA). Following ANOVA, pairwise comparisons were carried out using the post hoc Tukey honestly significant difference (HSD) test.

### Drosophila lifespan and diet manipulation analysis

The *w1118* parents were raised in vials containing standard food. To standardize the effects of parental age on offspring fitness, parents of experimental flies were of the same age (4∼5 days and reared at a constant density for at least two generations). To synchronize larval development, we allowed flies to lay eggs on yeast apple juice plates for 1 h, discarded the first batch of embryos, and then collected for another 4 h. Groups of 20∼25 embryos were put into vials containing standard food and allowed to develop until pupae eclosion. Newly eclosed flies were allowed to mature and mate for 48 h before the flies were briefly anesthetized with CO2 so that females and males could be separated for aging experiments. 100 adult flies were randomly allocated at a density of 20 flies per vial using 4 cohorts for each diet: standard food (ST), calorie restriction (CR), high glucose (HG), high sucrose (HS) and high fructose (HF). For the detail information of the food recipe, see Table S1. All eclosures were maintained at 25°C in a controlled light (12/12-h light/dark cycle) and humidity (>70%) environment. Flies were scored for survival daily and provided with fresh medium every 2 days. To minimize any density effects on mortality, two vials with cohorts were merged when the density of flies reached five or fewer individuals.

Eclosures were placed randomly in the incubator, and positions were rotated after each transfer to minimize the effects of microclimate. This process was followed until all flies were dead. Individual flies were censored from the results if they escaped during tipping or if got stuck in the food (non-age-related cause). For each diet treatment, median and maximum lifespan were calculated as half of deaths of the whole population and the longest lived 10% of individuals, respectively. Analysis of survival curve was performed using the Kaplan–Meier method, which takes account of censored data. The Kaplan–Meier estimate is the non-parametric maximum likelihood of survival at a given time-point. A plot of the Kaplan–Meier estimate is a series of horizontal steps of declining magnitude which, when a large enough sample is taken, approaches the true survival function for that population.

### D_2_O-labeling experiments

The lipogenic activity changes of wild type flies at different ages were labeled by transferring the 0-day, 10-day, 20-day, 30-day and 35-day adult flies to the 20% D2O labeled corresponding food conditions for 5 days, then the 5-day, 15-day, 25-day, 35-day, and 45-day aged flies were sacrificed, and their brains were dissected and subjected to Raman measurements and SRS imaging.

For investigating the lipogenic activity of the IIS related gene mutants, *chico*^*1*^/+ and *dfoxo*^*Δ94*^/*dfoxo*^*25*^ flies aged at 2 days (young) and 30 days (old) were labeled by 20% D2O labeled for 5 days, then the 7-day and 35-day aged flies were sacrificed, and their brains were dissected and subjected to Raman measurements and SRS imaging, respectively. The age paired *w*^*1118*^(*+/+*) flies treated in the same way were used as control.

For determining the lipogenic activity of glia-specific IIS manipulated transgenic flies. Groups of 15 *GliaGS* virgin females were collected to cross with 15 *UAS-InRCA, UAS-InRDN, UAS-Pten of UAS-dfoxo RNAi* males, respectively. All the crosses were maintained at 21°C on the standard food to allow the normal embryo and larvae development. 80 newly eclosed progenies (40 females and 40 males) with correct genotype were collected. They were allocated to 4 cohorts (20 flies of each) and transferred to the fresh food with RU-486 (Sigma, Cat #: 84371-65-3) added at a final concentration of 200 μM and raised in 25°C to allow the *Gal4-UAS* system function well. After 2 days maturation and mating, the 15 females and 15 males were separated and subjected to 5 day 20% D2O labeling experiments, then the brains from 7-day aged flies were dissected and measured by Raman or SRS imaging system. The rest of the flies (25 females and 25 males) were allowed to grow old to 30 days for D2O treatment and following experiments.

For the starvation assay, four groups of 2-/ 20-day old adult flies (15 flies per group) were transferred to different vials with 20% D2O labeled diets for 10 days labeling. Then the labeled (12-day and 30-day) flies were starved on 1% agar. At each time point (24 h, 48 h, 72 h after starvation), 5 files randomly selected from the four vials were sacrificed, and brains were dissected and subjected to SRS imaging for the C-D signal quantification.

The CD/CH ratios of LDs from three ROIs in each brain were measured by ImageJ software and then the quantifications from 5∼10 brains at each group were used to do statistical analysis.

### Brain LD staining

Brains were fixed overnight in 4% PFA in PBS in 4 °C, then washed 3 times with PBS. 1µg/mL BODIPY 493/503 (Invitrogen™, Cat# D3922) was used as final concentration to stain overnight in 4 °C. Following one rinse in PBS, tissues were rinsed and mounted in PBS for two photon fluorescent imaging at 800 nm wavelength.

### Statistical analysis

Kaplan-Meier Log-rank test was performed for survival assays. Statistical significance were tested by using student’s *t*-test, 1-way or 2-way ANOVA with a post hoc Tukey’s multiple comparison or ANOVA with a post hoc Dunnett’s comparison using GraphPad Prism software.

## Supplementary data

**Table S1.**
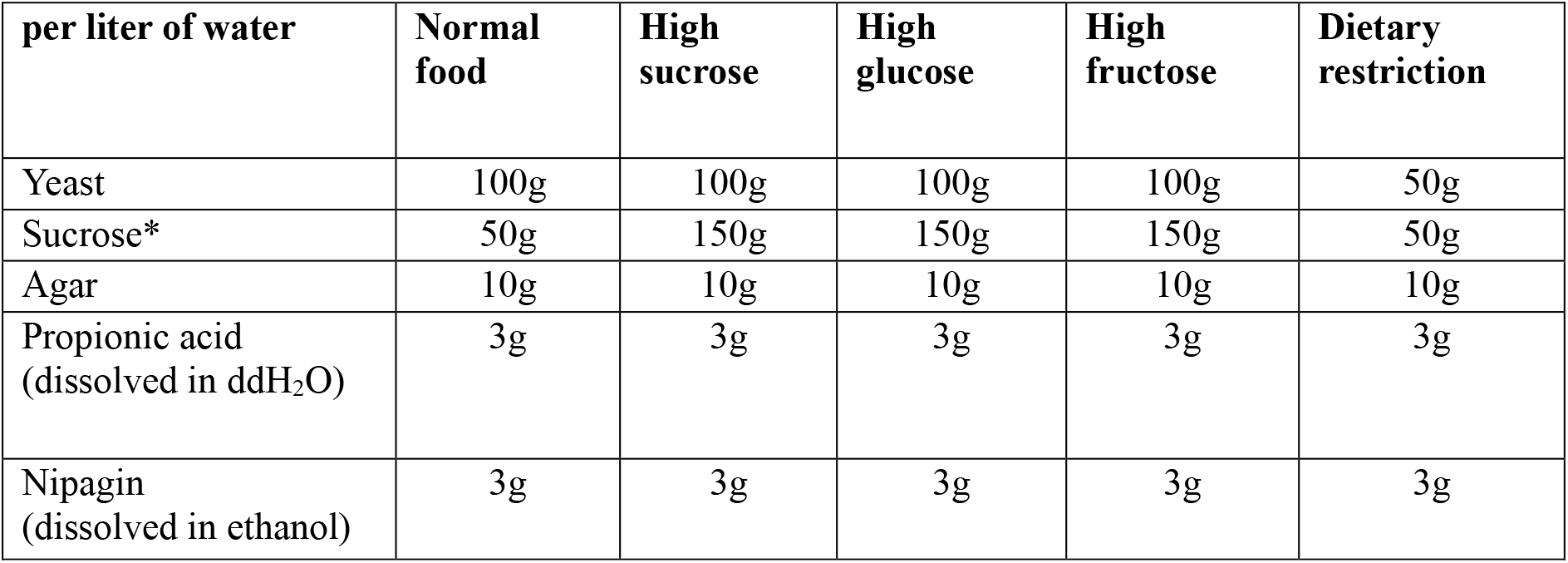
Food recipe for *Drosophila* diet treatment. **\*** For high glucose and high fructose diets, replace sucrose with glucose and fructose, respectively.

| per liter of water | Normal food | High sucrose | High glucose | High fructose | Dietary restriction |
| --- | --- | --- | --- | --- | --- |
| Yeast | 100g | 100g | 100g | 100g | 50g |
| Sucrose* | 50g | 150g | 150g | 150g | 50g |
| Agar | 10g | 10g | 10g | 10g | 10g |
| Propionic acid<br>(dissolved in ddH<sub>2</sub>O) | 3g | 3g | 3g | 3g | 3g |
| Nipagin<br>(dissolved in ethanol) | 3g | 3g | 3g | 3g | 3g |

**Figure S1.**
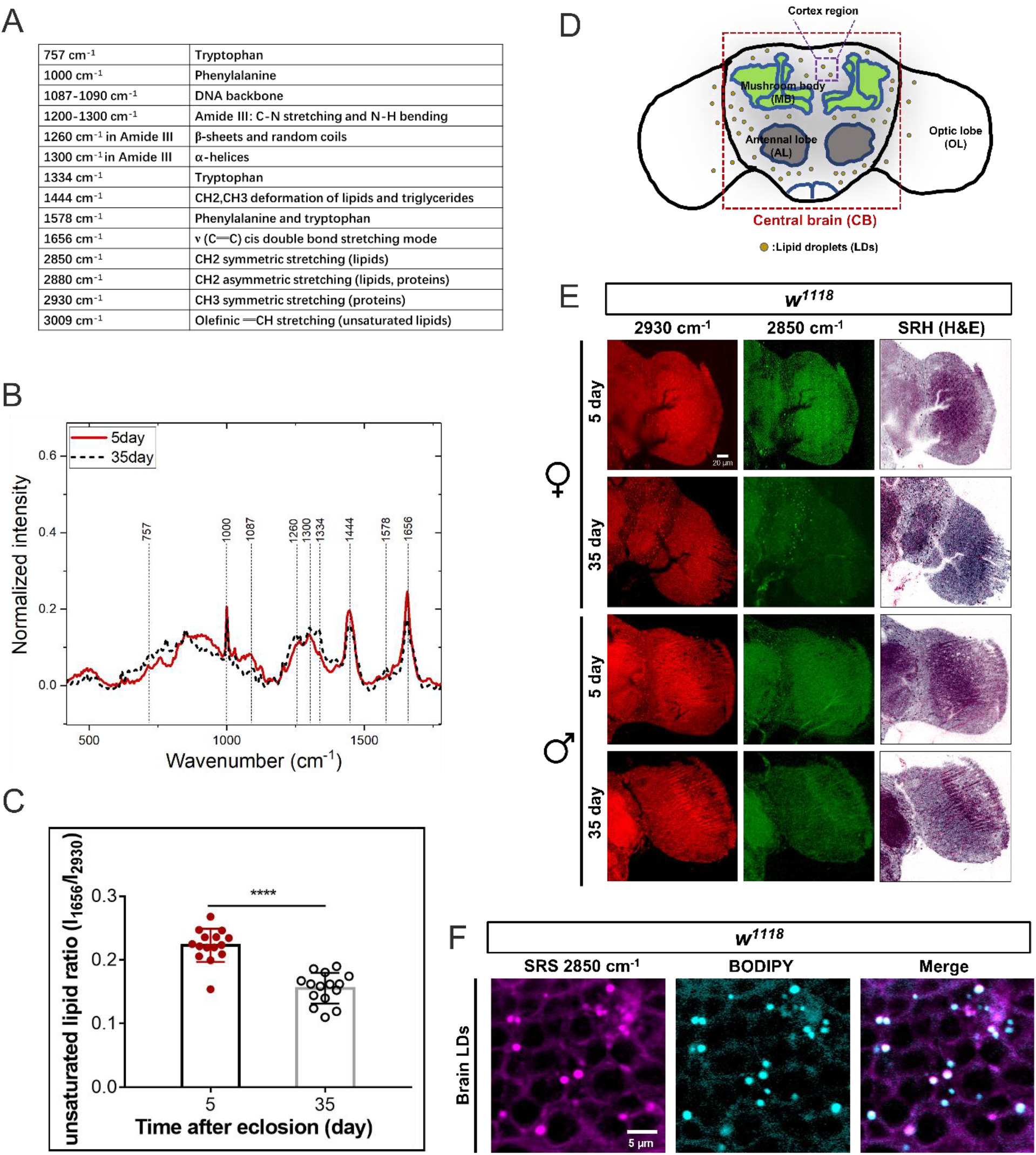
The lipid metabolic changes of *Drosophila* brain are visualized by SRS microscopy. (A) The Raman peaks in *Drosophila* brains are assigned to different vibrational modes. (B) The fingerprint regions of brain Raman spectra from 5-day and 35-day old flies were collected and plotted as Mean ± SD, n =15 brains for each age group. (C) The ratios of unsaturated lipids (1656cm^-1^) to proteins (2930 cm^-1^) level were quantified and plotted as Mean ± SD, n =15 brains for each age group. (D) A cartoon of *Drosophila* adult brain depicting the structures of central brain (CB), mushroom body (MB), antennal lobe (AL) and optical lobe (OL) as well as the cortex region. (E) The optic lobes from 5day and 35day flies were imaged from the proteins (2930 cm^-1^) and lipids (2850 cm^-1^) channels, from which signals were color-coded in red and green respectively. The stimulated Raman histology (SRH) H&E were generated from proteins and lipids channels. (F) The brain LDs imaged at SRS 2850 cm^-1^ channel are confirmed with BODIPY staining. Statistical significance was determined by using student’s *t*-test (C). *, p < 0.05; **, p < 0.01; ***, p < 0.001; ****, p < 0.0001; ns, non-significant difference.

**Figure S2.**
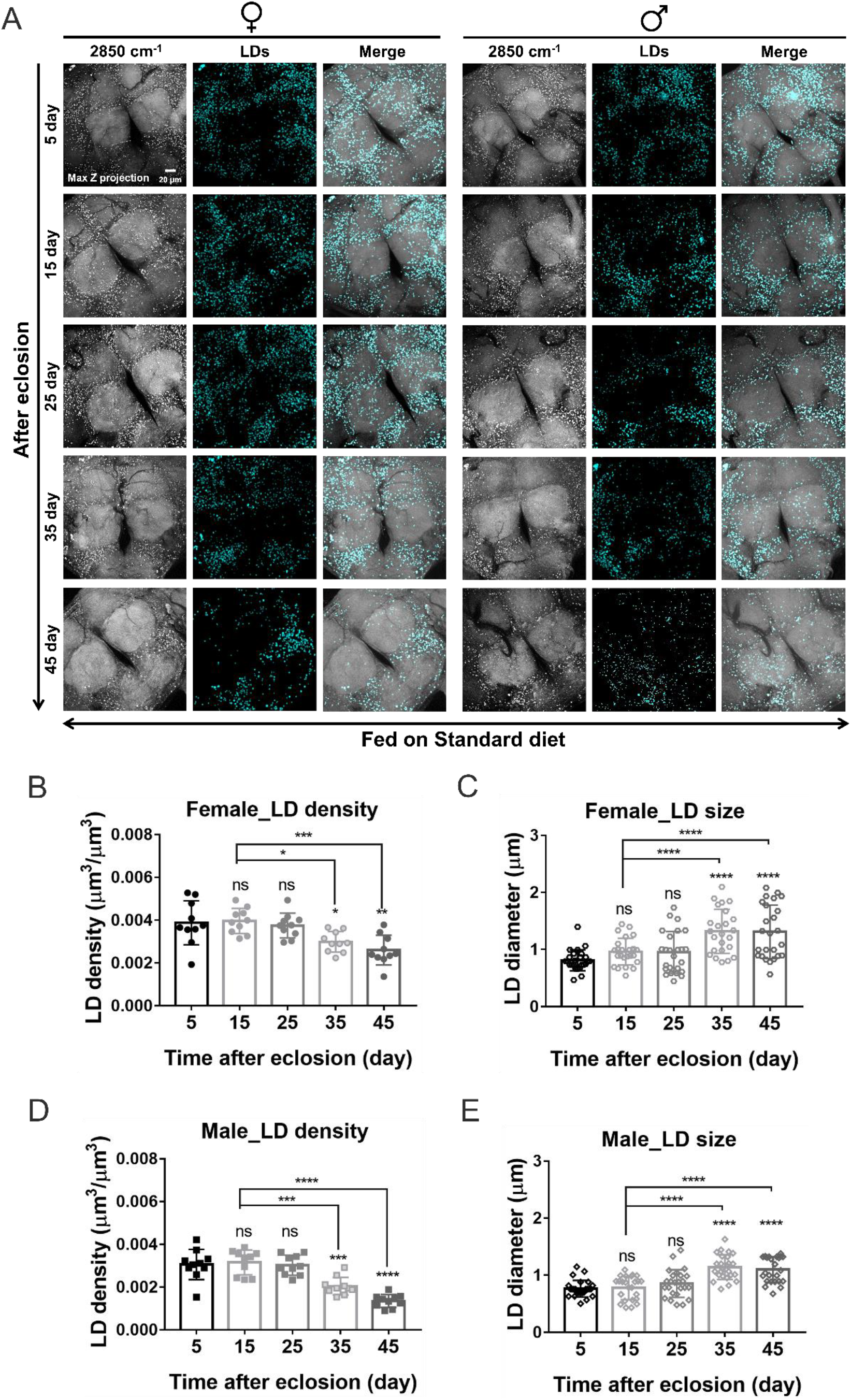
The LD density and size of *Drosophila* brain are changed during the aging process. (A) Max Z projected images of brain LDs from females and males at different ages were displayed. Scale bar, 10 µm. (B-E) The LD density (total LD volume to brain cortex volume) and LD size of different aged brains were quantified and presented as Mean ± SD (n = 8∼10 brains at each age group). Statistical significance was determined by using 1-way ANOVA (B-E). *, p < 0.05; **, p < 0.01; ***, p < 0.001; ****, p < 0.0001; ns, non-significant difference.

**Figure S3.**
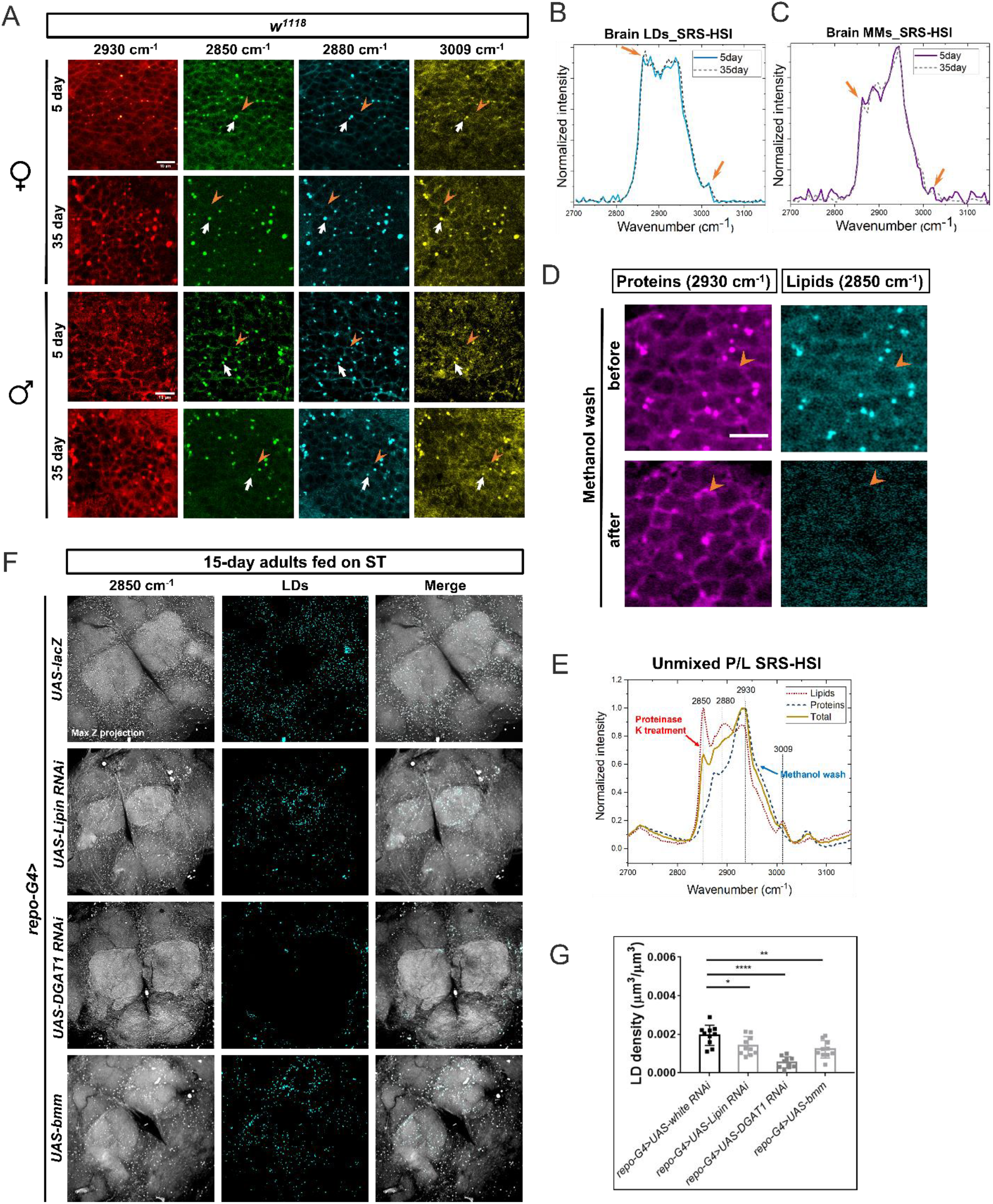
TAGs are the predominant constitution of glial LDs. (A) The cell body regions from 5-day and 35-day brain cortexes were imaged from the proteins (2930 cm^-1^), lipids (2850 cm^-1^), proteins and lipids mixture (2880 cm^-1^) and unsaturated lipids (3009 cm^-1^) channels. Signals were color-coded in red, green, cyan and yellow, respectively. White arrows, LDs; orange arrow heads, cell membranes. Scale bar: 10 µm. (B-C) The SRS-HSI were taken from 5- and 35-day old female brain LDs and membranes, and the spectra of CH regions from LDs (B) and membranes (C) were plotted. Total lipids peak (2850 cm^-1^) and unsaturated lipids peak (3009 cm^-1^) were pointed out by the orange arrows. (D) The cell body regions from 5-day adult brain were imaged from the proteins (2930 cm^-1^), lipids (2850 cm^-1^) channels before and after methanol treatment. Scale bar: 10 µm. (E) The spectra derived from SRS-HSI of untreated brains (yellow, solid line), proteinase K-treated ones (pure brain lipids, red dotted line) and methanol-treated brain membranes (pure brain proteins, blue dashed lines) showing the peaks under different conditions. (F) Max Z projected images of brain LDs with glial-specific gene knockdowns (*repo-G4>UAS-RNAi*) of TAG synthetic gene (*Lipin, DGAT1*) and overexpression of TAG catabolic gene (*bmm*) were displayed to compare with the control group (*repo-G4>UAS-lacZ*). (G) The LD density (ratios of total LD volume to brain cortex volume) in (F) were quantified and presented as Mean ± SD (n =5∼8 brains). Statistical significance was determined by using Student’s *t*-test (D, E) or 1-way ANOVA (G). *, p < 0.05; **, p < 0.01; ***, p < 0.001; ****, p < 0.0001; ns, non-significant difference.

**Figure S4.**
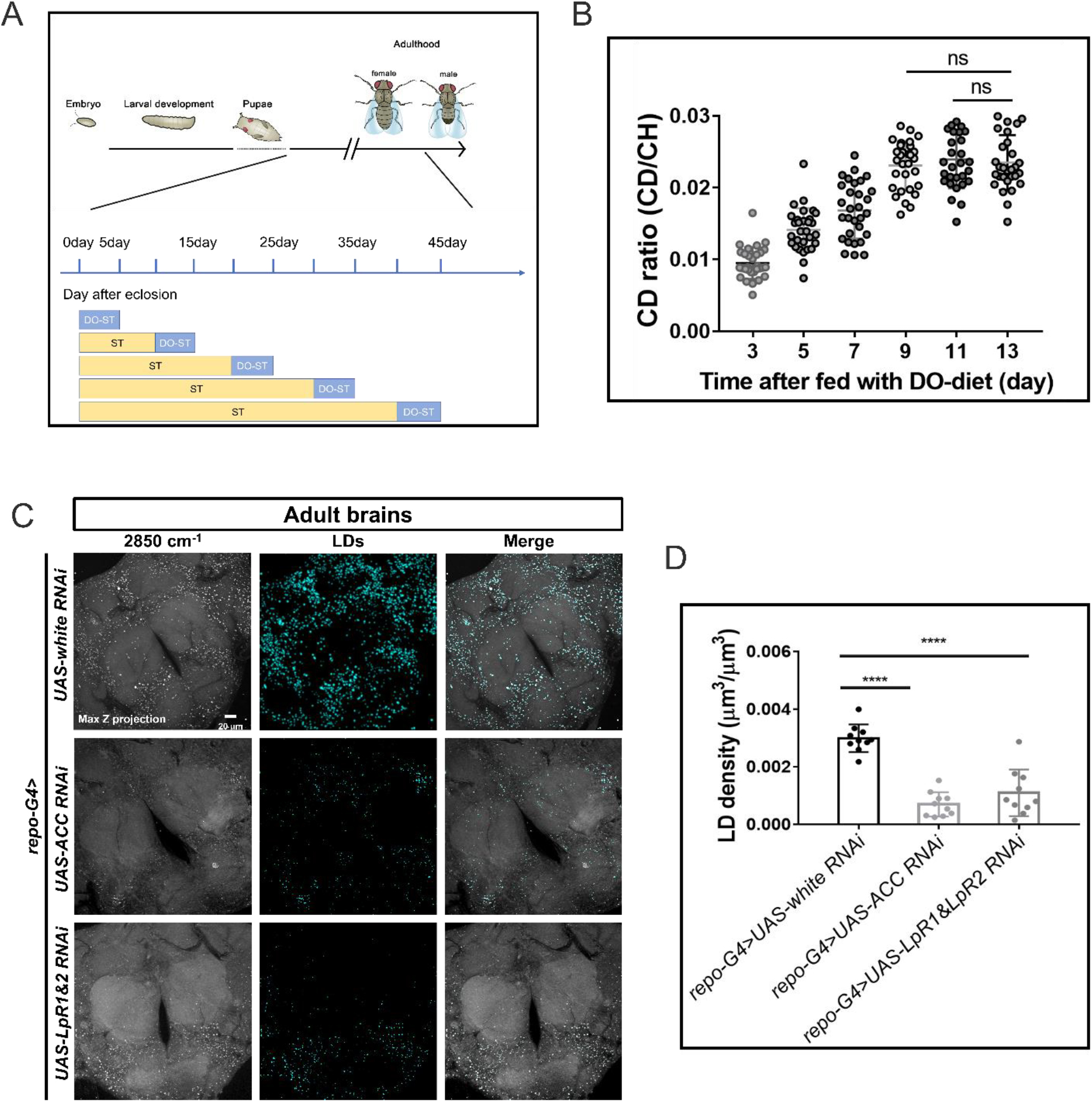
The *de novo* biosynthesis and uptaking of lipids are both contributing to the brain lipid storage. (A) A scheme showing D2O diet (DO-ST, D2O labeled standard diet) treatment of adult flies during aging. (B) The deuterium incorporation of brain LDs reaches saturation after 9∼10 days’ 20% D2O diet treatment. (n =5 brains at each time point). (C) Max Z projected images of brain LDs with glial-specific gene knockdowns (*repo>RNAi*) of fatty acid synthetic gene (*ACC*) and the genes encoding lipid uptaking receptor (*LpR1&LpR2*) were displayed to compare with the control group (*repo>UAS-white RNAi*). (D) The LD density ratios (total LD volume to brain cortex volume) of the brains with the designated genotypes in (C) were quantified and presented as Mean ± SD (n =5 brains in each group). Statistical significance was determined by using 1-way ANOVA. *, p < 0.05; **, p < 0.01; ***, p < 0.001; ****, p < 0.0001; ns, non-significant difference.

**Figure S5.**
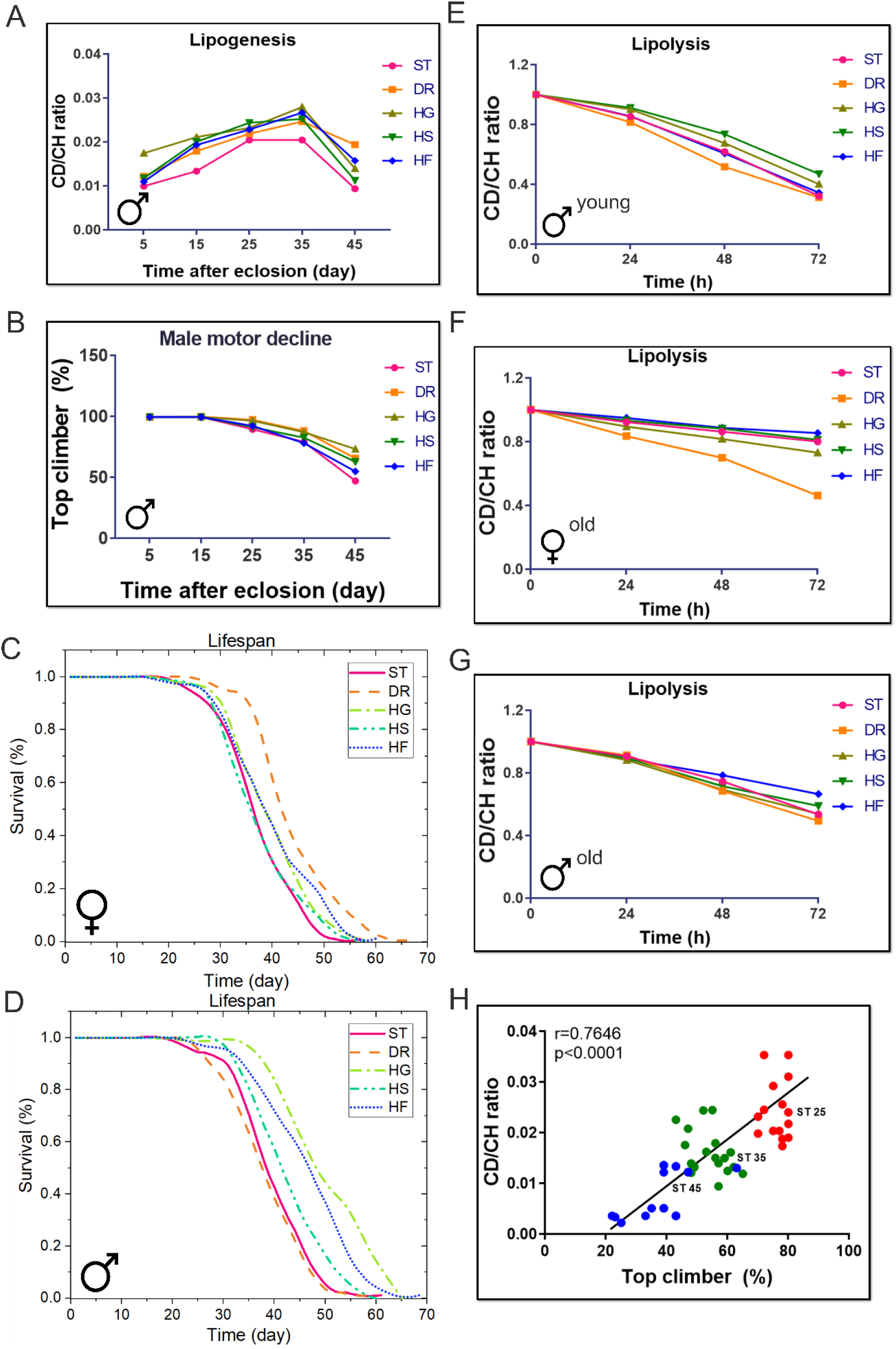
The LD metabolic activity of aging brains under diet treatments are correlated with motor activity and lifespan. (A) The brain LDs from 5, 15, 25, 35 and 45-day old males fed with 20% D2O labeled standard diet (ST), high sugar (glucose, fructose and sucrose) diets (HSDs) and dietary restriction (DR) for 5 days were imaged from the CD lipids (2143 cm^-1^), CH lipids (2850 cm^-1^), and the ratio of newly synthesized lipids to total lipids during 5-days’ DO-labeling were quantified as CD ratios (CD/CH) and plotted as Mean ± SD. n =15 ROIs from 5∼8 brains. (B) The motor activities of 5, 15, 25, 35 and 45-day old males fed with standard diet (ST), high sugar (3xglucose, fructose and sucrose) diets (HSDs) were evaluated and plotted as Mean ± SD. N = 40 files per age group. (C) and (D) The lifespans of flies fed with standard diet (ST), high sugar (3xglucose, fructose and sucrose) diets (HSDs) were examined and plotted as in (C, females) and (D, males). (n =120 flies at each group). (E-G) The time-course changes of CD-lipid mobilization in brain LDs over 72-hour starvation of flies pretreated with different diets were quantified and plotted as Mean ± SD in (E, young male), (F, old female) and (G, old male). n =15 ROIs from 3∼5 brains in each group. (H) A linear positive correlation between brain lipid synthesis and motor ability during aging process. n = 10∼15 for each group, Pearson analysis, r = 0.7646, p < 0.0001.

**Figure S6.**
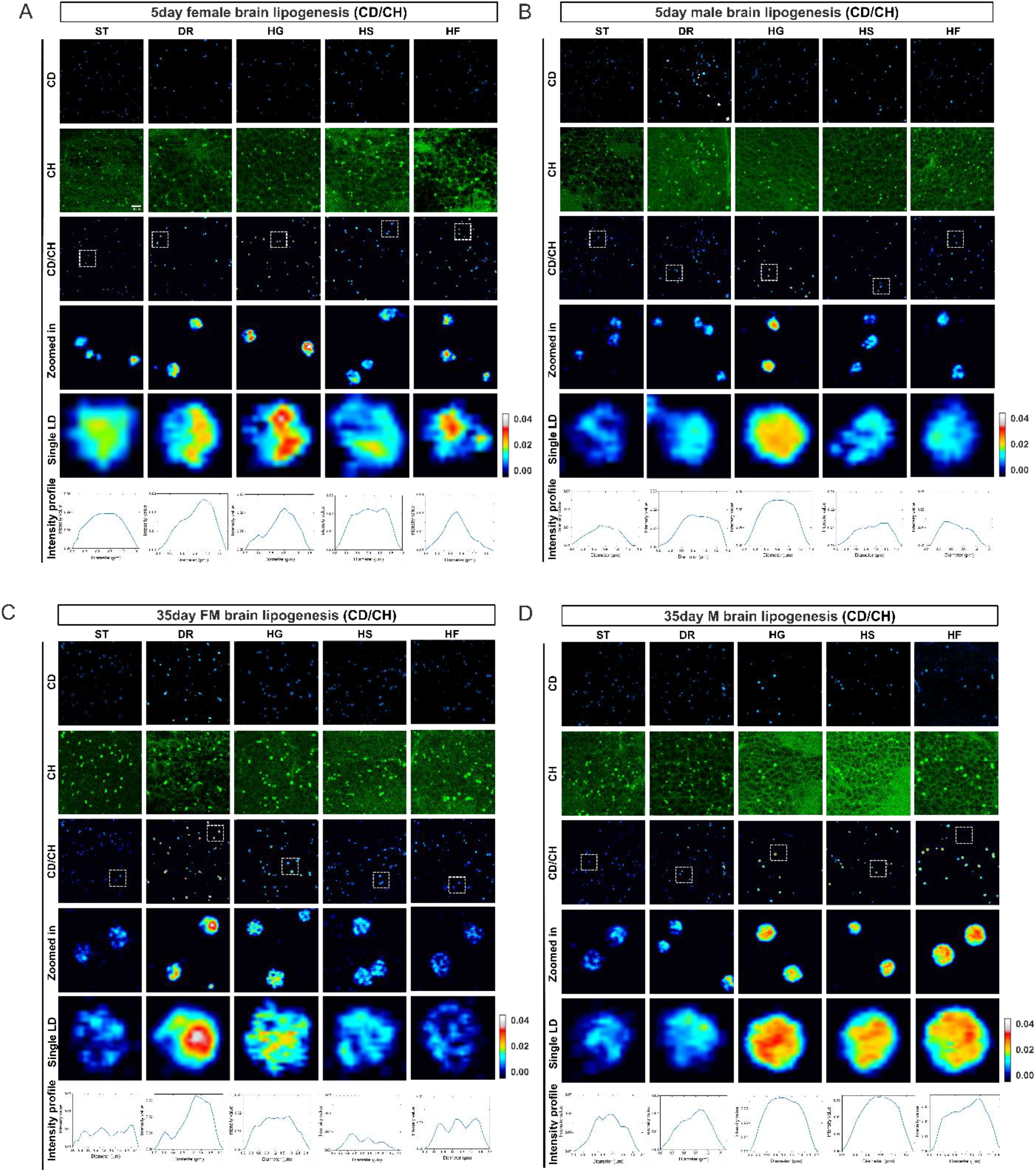
The lipogenic activities of aging brain LDs are modulated by diet treatments. (A-D) The brain LDs from 5, 15, 25, 35 and 45-day old flies fed with 20% D2O labeled standard diet (ST), high sugar (glucose, fructose and sucrose) diets (HSDs) and dietary restriction (DR) for 5 days were imaged from the CD lipids (2143 cm^-1^), CH lipids (2850 cm^-1^), and the ratio of newly synthesized lipids to total lipids during 5-days’ DO-labeling were quantified as CD ratios (CD/CH) and displayed in (A, 5-day females), (B, 5-day males), (C, 35-day females), (D, 35-day males), respectively. Scale bar: 2µm.

**Figure S7.**
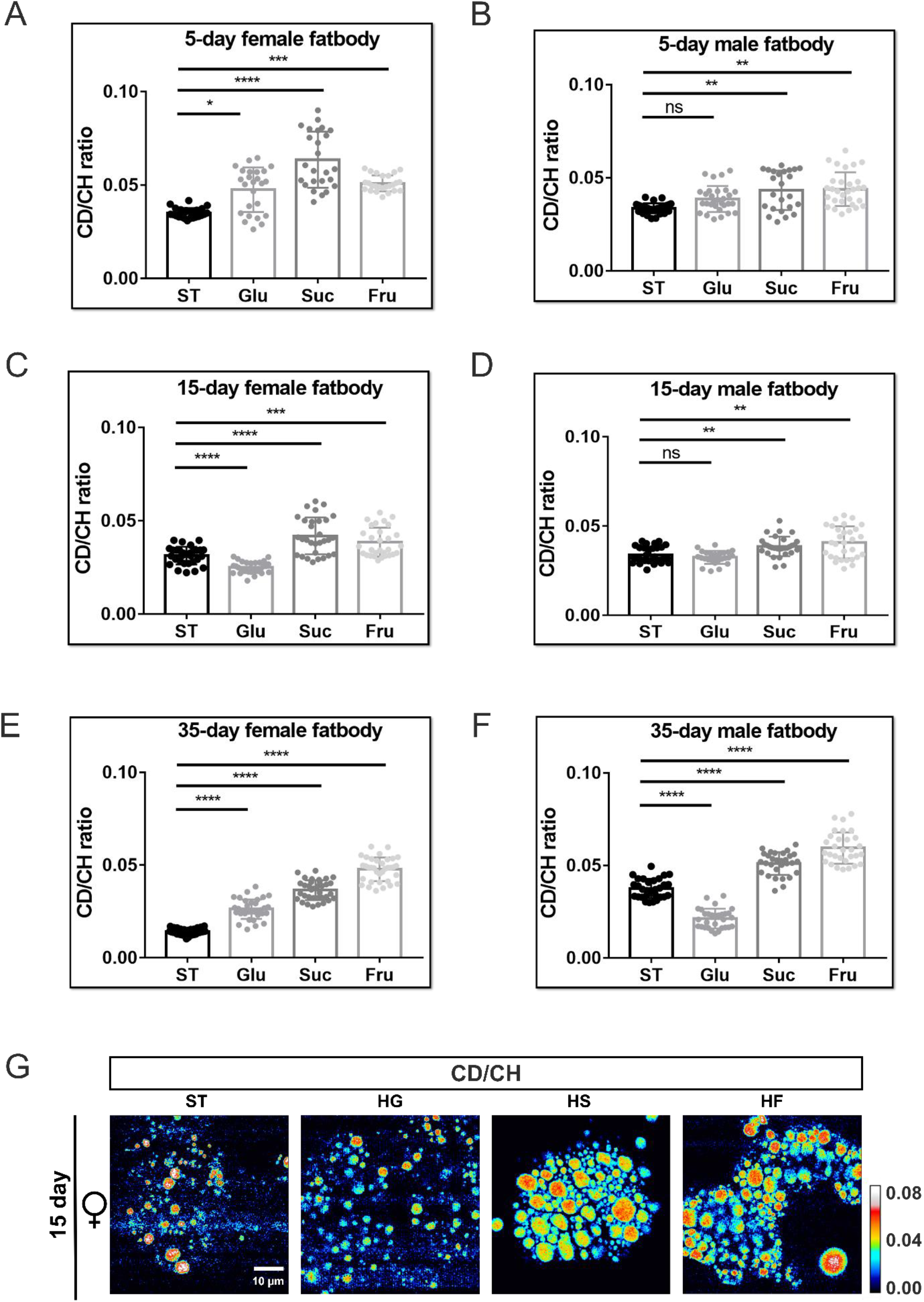
The lipogenic activities of LDs in fat bodies during aging are differently modulated by dietary sugars. (A-D) The LDs from 5, 15, and 35-day old flies fed with 20% D2O labeled standard diet (ST), high sugar (glucose, fructose and sucrose) diets (HSDs) for 5 days were imaged from the CD lipids (2143 cm^-1^), CH lipids (2850 cm^-1^), and the ratio of newly synthesized lipids to total lipids during 5-days’ DO-labeling were quantified as CD ratios (CD/CH) and displayed in females (A, C, E,), males (B, D, F) respectively. (n =25 ROIs from 5∼8 fat bodies). The representative radiometric images for lipid biosynthesis of 15-day female fat body LDs were displayed in (G), respectively. Scale bar: 10µm.

**Figure S8.**
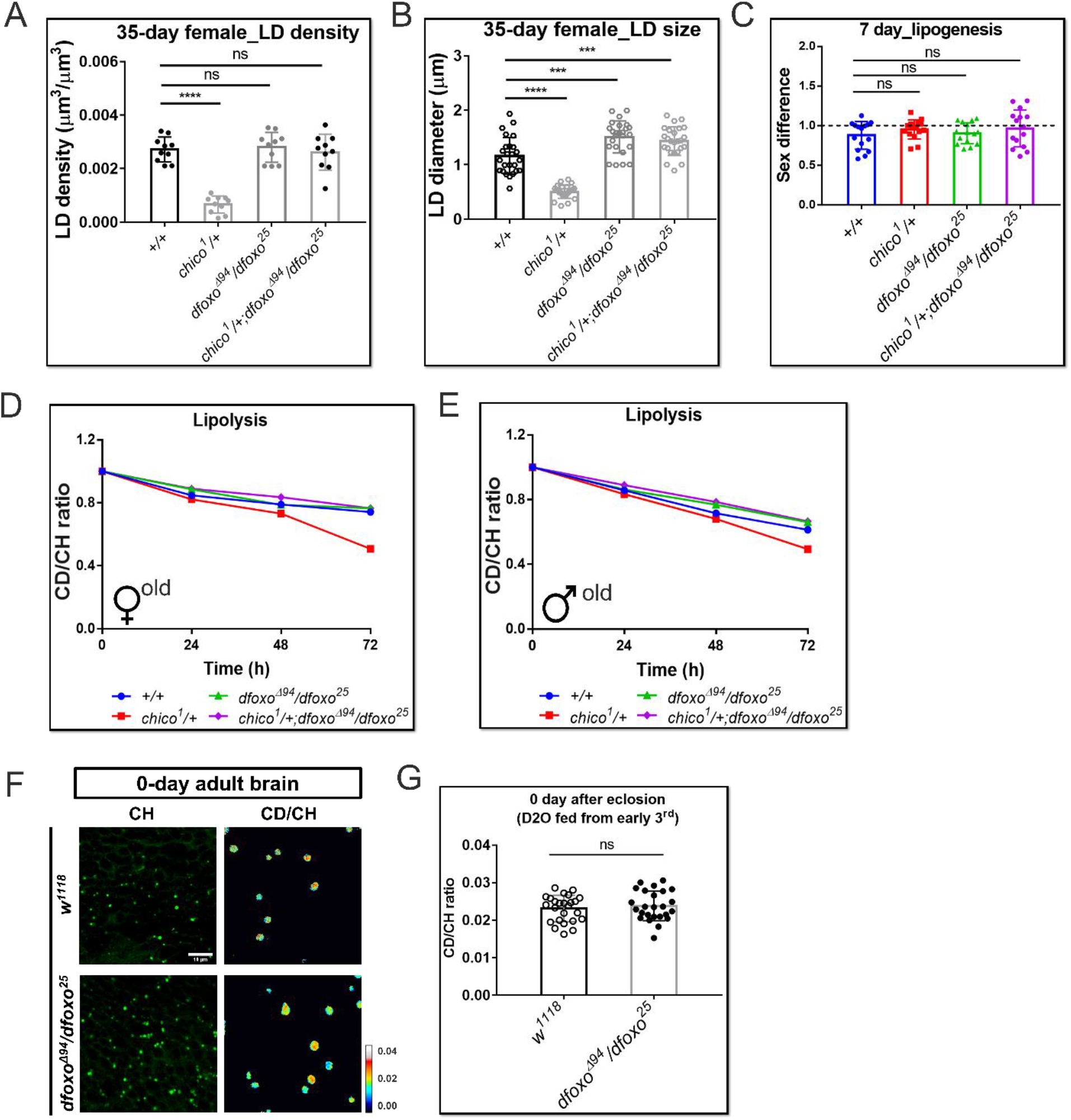
The lipid metabolic changes of genetic mutant brains in IIS pathway. (A-B) The LD density ratios (total LD volume to brain cortex volume) and LD sizes were quantified and presented as Mean ± SD in (A) and (B), respectively. N =5∼8 brains for each group. (C) The sex differences of lipogenesis in 7-day old IIS mutants were determined. (D-E) The time-course changes of CD-lipid mobilization after 0, 24, 48, 72-hour starvation of young and old brain LDs were quantified and plotted as Mean ± SD in (D, old female) and (E, old male), n =15 ROIs from 3∼5 brains. (F-G) The radiometric images for lipid biosynthesis of 0-day brain LDs from *dfoxo*^*Δ94*^*/dfoxo*^*25*^ and *w*^*1118*^ were displayed in (F) and quantified in (G). n =15 ROIs from 3∼5 brains.

**Figure S9.**
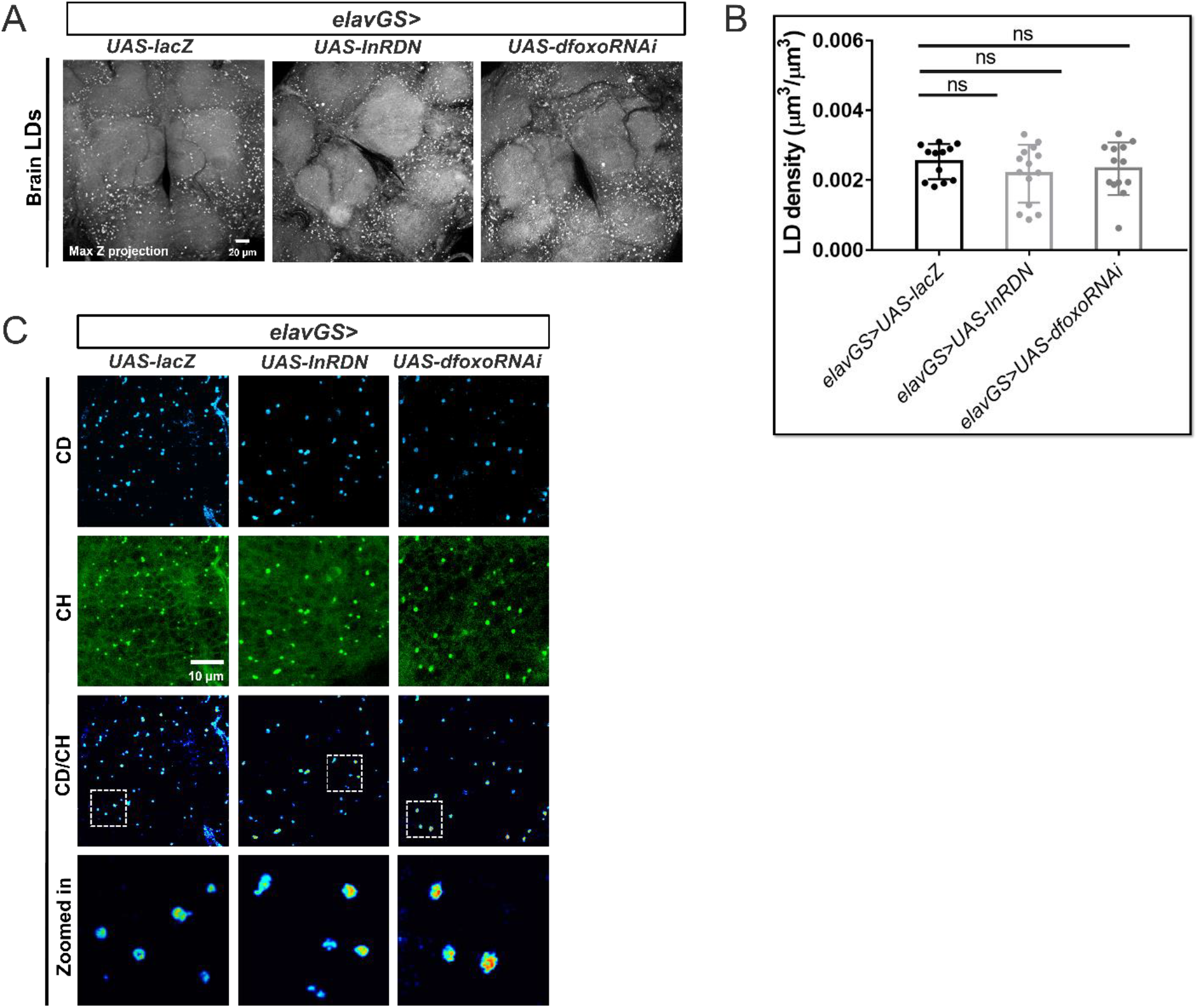
Neuron-specific manipulation of IIS pathway may not significantly affect the brain lipid metabolism. (A) Max Z projected images of brain LDs with neuron-specific gene manipulations of adult flies (*elavGS>UAS-InRDN* and *UAS-dfoxoRNAi*) are compared with that of *elavGS>UAS-lacZ* flies. (B) The LD density ratios (total LD volume to brain cortex volume) of the brains from the designated genotypes in (A) were quantified and presented as Mean ± SD (n = 5 brains for each group). (C) The images of brain LDs from genetic manipulated IIS flies at 10-day old flies treated with 20% D2O labeled standard diet (ST) for 5 days were displayed as green (CH, total lipids), cyan (CD, newly synthesized lipids), and royal (CD/CH, newly synthesized lipids to total lipids) respectively. The representative LDs inside the dashed box were magnified to show the details, Scale bar: 10µm.

**Figure S10.**
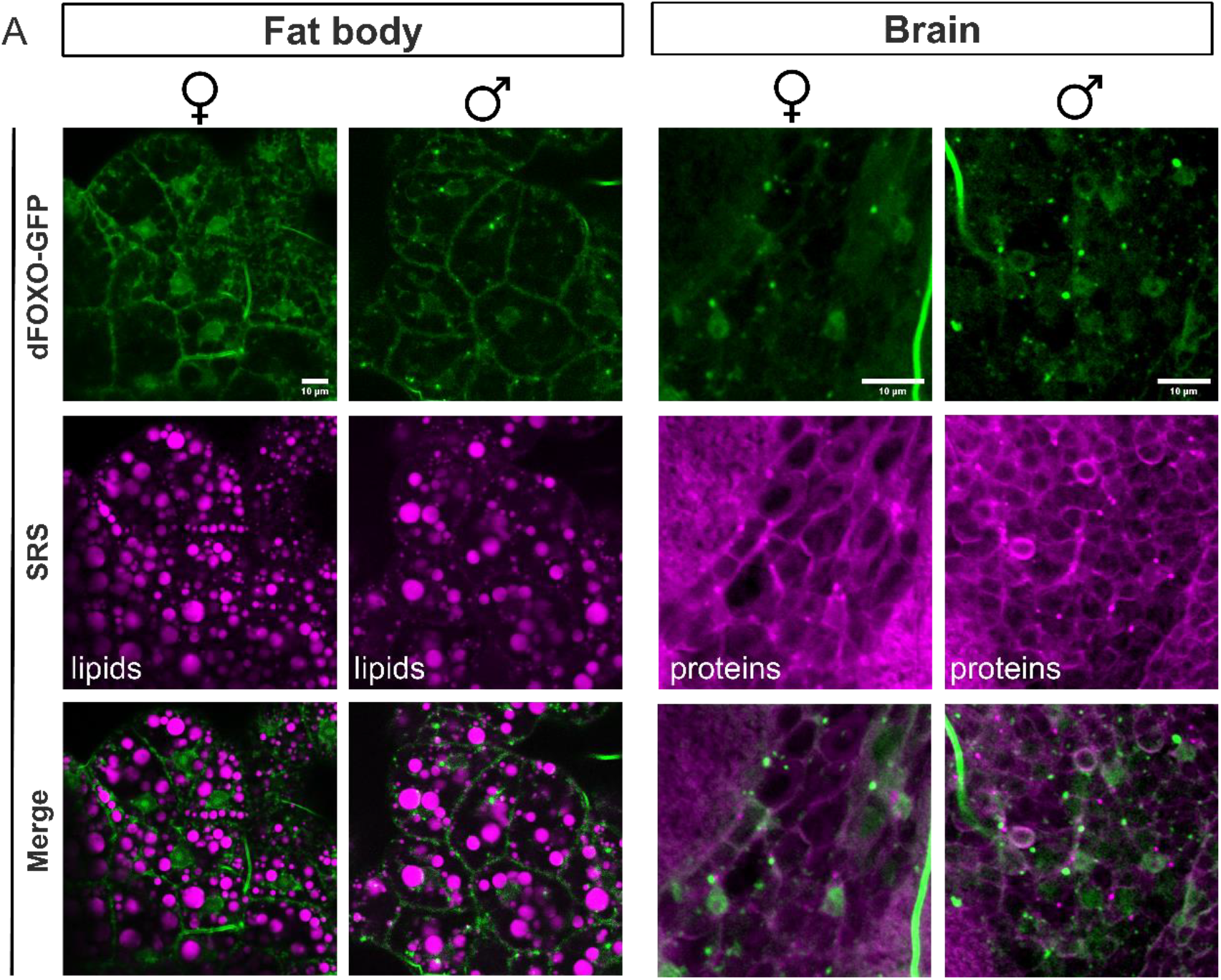
The IIS activities show sexual-dependent differences in adult fat bodies but may not in brains. (A) The fluorescent images of dFOXO-GFP expression in fat bodies and brains were taken from 10-day old female and male flies and were displayed as green channel, the SRS lipids in fat bodies and proteins in brains is assigned to magenta, respectively. The overlapped images as shown are generated based on fluorescent and SRS signal. Scale bar: 10µm.

